# Molecular mechanisms of heavy metal adaptation of an extremophilic red alga *Cyanidioschyzon merolae*

**DOI:** 10.1101/2023.02.24.529964

**Authors:** Francesca Marchetto, Sergio Santaeufemia, Magdalena Lebiedzińska-Arciszewska, Małgorzata A. Śliwińska, Magdalena Pich, Eliza Kurek, Aleksandra Naziębło, Marcin Strawski, Daniel Solymosi, Marek Szklarczyk, Ewa Bulska, Jędrzej Szymański, Małgorzata Wierzbicka, Yagut Allahverdiyeva-Rinne, Mariusz R. Więckowski, Joanna Kargul

**Author notes:** Address for correspondence. The author responsible for the distribution of materials integral to the findings presented in this article following the policy described in the Instructions for Authors (www.plantphysiol.org) is Joanna Kargul. F.M., S.S., M.L.A., M.A.Ś., M.S. and MP generated and processed the data and prepared the figures; J.K., F.M., M.R.W. and S.S. designed the experiments, analyzed and interpreted the data; M.Sz., E.B., Y.A.R., J.S., M.R.W. analyzed and interpreted the data; J.K., F.M. and S.S. co-wrote the article; J.K. conceived, conceptualized,coordinated the study and obtained funding. J.K., F.M., and S.S. gratefully acknowledge the support from the Polish National Science Centre (OPUS 17 grant no. 2019/33/B/NZ3/01870 to J.K. and MINIATURA 5 grant no. 2021/05/X/NZ2/01516 to S.S.S.).

## Abstract

The order of Cyanidiales comprise seven acido-thermophilic red microalgal species thriving in hot springs of volcanic origin characterized by extremely low pH, moderately high temperatures and the presence of elevated concentrations of sulphites and heavy metals that are prohibitive for most other organisms. Little is known about the molecular mechanisms of Cyanidiales long-term adaptation to such hostile environments, in particular to heavy metals, yet elucidation of these processes is important for understanding the evolution of the metabolic pathways underlying heavy metal detoxification for developing rational strategies for heavy metal bioremediation. Here, we investigated the long-term adaptive responses of *Cyanidioschyzon merolae* cells, a member of Cyanidiales, to extremely high nickel concentrations. Through complementary approaches based on physiological, microscopic and elemental analyses we dissect several molecular mechanisms underlying the long-term adaptation of this model extremophilic microalga to high Ni exposure. These include: (i) extrusion of Ni from the cells and lack of significant Ni accumulation inside the cells; (ii) maintenance of efficient photoprotective responses including non-photochemical quenching and state transitions; (iii) dynamic remodeling of the chloroplast ultrastructure such as formation of metabolically active prolamellar bodies and plastoglobuli together with loosening of the thylakoid membranes; (iv) activation of ROS amelioration metabolic pathways; and (v) preservation of the efficient respiratory chain functionality. All the dynamically regulated processes identified in this study underlie the remarkable adaptability of *C. merolae* to extremely high Ni levels that exceed by several orders of magnitude the levels of this heavy metal found in the natural environment of this extremophile.

## 1. Introduction

The vast majority of environmental conditions on Earth are hospitable to most organisms. However, there are places where conditions remain similar to those observed in the early onset of life. The adverse conditions of such environments, where liquid water or energy supplementation are scarce, are detrimental to the vast majority of organisms, with values that are far from the normal standards for living organisms to prosper. Nevertheless, extremophilic microorganisms can cope with these hostile environments naturally by effectively adapting to the extreme external conditions (Aguilera, 2013; Seckbach, 2010). Dissecting the intricacies of metabolic regulation of homeostasis in extremophilic microorganisms is pivotal for understanding the limits at which life can flourish. Since the conditions in which extremophiles are capable of survival are similar to those in which the first cells evolved, understanding the molecular mechanisms that underpin life in the extreme environments can shed light on the origin of life.

Different species of extremophiles inhabit environments that are hostile for most forms of life. Among those, particularly fascinating is the rhodophytan order Cyanidiales that comprises the extremophilic red microalgae thriving in hot springs of volcanic origin. The abiotic conditions of such environments include very low pH (0.05-4), moderately high temperatures (40-57°C) and the presence of elevated concentrations of sulphites and heavy metals that are prohibitive for virtually all other forms of life. The order comprises a mere seven species of unicelular algae (*Cyanidium caldarium, Cyanidioschyzon merolae, Galdieria partita, Galdieria daedala, Galdieria maxima, Galdieria phlegrea* and *Galdiera sulphuraria)* (Ciniglia et al., 2004) which form the majority of biomass under these challenging environmental conditions. The heavy metal concentrations associated with these ecosystems are significantly higher than those permitted by the international toxicity regulating bodies. As an example, concentrations of 0.058, 0.023, 0.571 ppm of Ni, Cd and Cr, respectively, have been found in these environments (Shakhatreh et al., 2017); which is 3-11-fold higher than the limits set by the World Health Organization corresponding to 0.02, 0.003 and 0.05 ppm for Ni, Cd and Cr, respectively (Luca et al., 1978; Carr and Neary, 2008).

Among them, one of the most studied species is a model extremophile, *Cyanidioschyzon merolae*, a unicellular thermo-acidophilic red microalga, isolated from volcanic hot springs in Campi Flegrei, Naples, Italy (Luca et al., 1978). This species has attracted a great deal of interest because of its evolutionary position at the root of the red algal lineage, which would correspond to a basal group of eukaryotes (Miyagishima et al., 2017). On the other hand, *C. merolae* has been considered an evolutionary link between prokaryotic cyanobacteria and photosynthetic eukaryotes, as manifested e.g. through the hybrid character of the structure and function of the photosynthetic apparatus. Thus, the photosystem II (PSII) complex (water splitting enzyme) contains highly conserved core/reaction center subunits, prokaryotic oxygen evolving complex subunits (PsbV and PsbU) in addition of the conserved PsbO and a fourth unique PsbQ’ subunit. The light harvesting antenna of PSII is formed by the cyanobacterial-like phycobilisome (PBS) complex, containing c-phycocyanin and allophycocyanin as the main constituents. The photosystem I (PSI) complex is of a eukaryotic type, containing a crescent-like LHCI complex associated with the core complex on the PsaF/PsaJ side. Both photosystems (PS) contain exclusively chlorophyll *a* (Chl *a*) in contrast to the much richer composition of Chl isoforms in other phototrophs. The cell structure of *C. merolae* is very simple, with only a single nucleus, a single mitochondrion and a single chloroplast. Noteworthy, *C. merolae*, in contrast to other Cyanidiales, does not contain the cell wall or typical vacuoles, although electron-dense bodies of single membrane-bound, vacuole-like organelles rich in polyphosphate have been identified (Yagisawa et al., 2007).

All the three genomes are fully sequenced, making this microalga an attractive as a model system to dissect the evolution and molecular components of the signaling pathways related to the fundamental cellular processes including oxygenic photosynthesis (Busch et al., 2010; Krupnik et al., 2013; Nikolova et al., 2017; Tian et al., 2017; Haniewicz et al., 2018; Langley et al., 2018; Antoshvili et al., 2019; Abram et al., 2020; Chang et al., 2020),as well as cell cycle, cell division, protein and lipid homeostasis and circadian rhythms (Kuroiwa, 1998; Kuroiwa et al., 1998; Imamura et al., 2015; Miyagishima and Tanaka, 2021).

Considerable research has focused on the application of extremophilic microalgae in biotechnology (Varshney et al., 2015), including metal biomining and recycling (Cho et al., 2020) and biofuel production (Cheng et al., 2019; Dandamudi et al., 2020; Lang et al., 2020) due to their unique metabolic traits and high enzymatic activity and stability. Yet the in-depth knowledge of the physiological responses and molecular components associated with the long-term adaptation of Cyanidiales to severe stressors such as high concentrations of heavy metals, high temperatures, high light and extremely acidic pH remains limited. In particular, despite a plethora of evidence pointing towards tolerance of Cyanidiales to elevated levels of heavy metals (Varshney et al., 2015), little is known about the precise molecular mechanisms underlying the heavy metal defense/detoxification strategies at the cellular level or the effects of heavy metals on the structure and function of the photosynthetic apparatus in these fascinating ‘living fossils’ that drive primary biomass production in the extreme environments of volcanic origin.

Elevated levels of heavy metals, such as iron, copper, cadmium, mercury, nickel, lead, and arsenic, can induce generation of reactive radicals and cause cellular damage via depletion of enzyme activities, through lipid peroxidation, and reaction with nuclear proteins and DNA. Much of the knowledge on heavy metal detoxification pathways comes from numerous studies in higher plants and remains largely unknown for extremophilic red microalgae. At the cellular level, the defense mechanisms to a metal ion can be either exclusion of the metal from the cell, the uptake and modification of the metal to a less toxic form, followed by the metal’s extrusion from the cell, or by internal sequestration mainly in vacuoles (Hall and Williams, 2003). These processes are a result of a coordinated network of biochemical processes, which increase the cell’s ability to maintain homeostasis, and minimize oxidative stress resulting from the production of reactive oxygen species (ROS). Metal intoxication increases ROS levels as they play a significant role in many ROS-producing mechanisms, including the Haber–Weiss cycle, Fenton’s reactions, and disruption of the photosynthetic electron chain leading to superoxide and singlet oxygen production and reduction of the glutathione pool (Pinto et al., 2003). ROS also act as signaling molecules that induce the production of a network of antioxidants, antioxidant enzymes, and other stress related molecules (Shcolnick and Keren, 2006).

The main objective of this study was to provide comprehensive insight into the long-term adaptation mechanisms of *C. merolae* cells to high concentrations of heavy metals, specifically nickel. By using a combination of advanced spectroscopic, microscopic and elemental analyses we demonstrate the remarkable adaptability of *C. merolae* to extremely high (mM) concentrations of this heavy metal, with the concomitant preservation of high cell viability, high photosynthetic performance, including efficient photoprotective mechanisms, and triggering reorganization of the thylakoid architecture, formation of new sub-organelles (plastoglobules and prolamellar bodies) and activation of reactive oxygen species amelioration pathways.

## 2. Results and Discussion

### 2.1. Growth and viability of C. merolae cells in presence of high concentrations of nickel

The main aim of this study was to gain insight into the mechanisms of long-term adaptation of *C. merolae* cells to nickel, in particular examine the associated functional changes of the photosynthetic apparatus that provides the majority of metabolic energy required for maintenance of cellular homeostasis under adverse conditions such as metal stress. Nickel is an essential element for most organisms, and specifically it is a micronutrient required for microalgae cell growth (Muyssen et al., 2011). However, in higher plants and mesophilic algae high concentrations of this heavy metal negatively affect the cell growth and division due to the inhibition of enzymatic activity including the function of the components of the photosynthetic apparatus (Monteiro et al., 2012).

As the starting point of our study, we examined the growth curves of *C. merolae* cells exposed to 1, 3, 6- and 10-mM Ni concentrations (58.6, 175.8, 351.6 and 586 ppm, respectively) for up to 15 days (Fig. 1). The Ni concentrations applied in our study are 1-2 orders of magnitude higher than those reported for the majority of aquatic environments. For instance, Ni concentrations of 3.88 x 10^-6^ – 10.88 x 10^-6^ mM or 3.41 x 10^-5^ – 17.04 x 10^-5^ mM have been found in seawater (Bruland et al., 1979) and freshwater (Quality et al., 1981) environments, respectively. On the other hand, the Ni concentration in the hot volcanic springs from which *C. merolae* was isolated is in the range of 0.2-9.1 ppm, which corresponds to 0.003-0.155 mM Ni (Piochi et al., 2019). Therefore, it is likely that this extremophilic microalga may have evolved efficient adaptation mechanisms that allow it to thrive in the Ni-rich environments, including the anthropogenically polluted environments.

**Figure 1.**
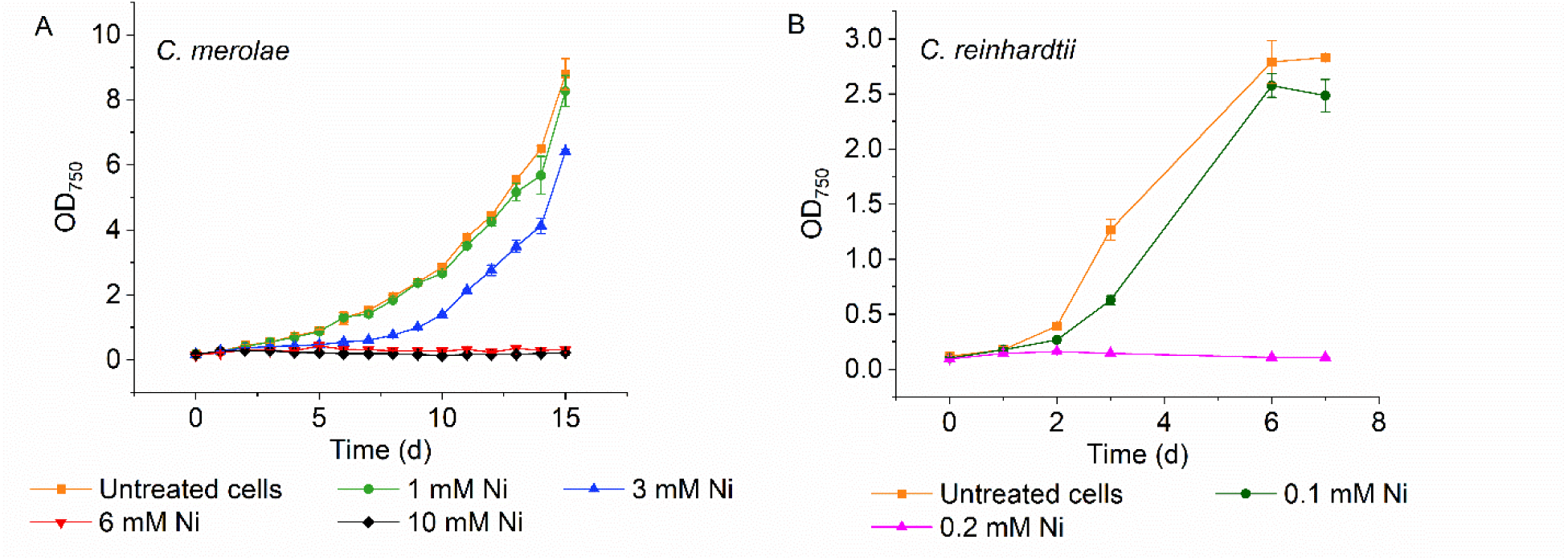
Growth curves of *C. merolae* (A) and *C. reinhardtii* (B) cell suspensions exposed to various nickel concentrations. Cell growth was assessed by measuring OD_750_ values of cell suspensions during up to 15-day exposure to 0.1-10 mM Ni. Data represents the mean ± SD of two independent biological replicas (n=2).

We observed that up to 1 mM Ni in the growth medium, no significant decrease in cell growth occurred compared to the control (p < 0.05 for day 5, 10 and 15; Fig. 1), which confirms the adaptability of *C. merolae* to high Ni concentration in the millimolar range. On the other hand, cultures exposed to Ni concentrations higher than 1 mM, i.e., 3, 6 and 10 mM, showed gradual inhibition of cell growth from day 2 of the treatment compared to the untreated cells (p < 0.0001). This inhibitory effect was more severe in the case of 6 mM and 10 mM Ni treatment than in the presence of 3 mM Ni, whereby the optical density at 750 nm (OD_750_) significantly decreased after 5 days of Ni exposure. However, in the case of 3 mM Ni, there was an observable cell growth in the first 3 days, then a lag phase until day 5 followed by logarithmic growth from day 6 until the last day of the experiment (day 15), albeit with slower growth rate compared to the control and cells exposed to 1 mM Ni (p < 0.01). The observed cell growth recovery at 3 mM Ni indicates triggering the adaptation mechanisms in *C. merolae* cells that allow these cells to thrive in the presence of such highly toxic concentrations of Ni. The fitting of the growth data of *C. merolae* in presence of different Ni concentrations allowed us to determine the Ni concentration at which the cell growth (monitored at OD_750_) is reduced by 50% (IC_50_) after 15 days of Ni treatment, which corresponds to 3.94 µM Ni (Fig. S1).

These results were compared with those obtained with the mesophilic green microalga *Chlamydomonas reinhardtii*, which inhabits environments with neutral pH and temperatures around 20-32℃. Growth of *C. reinhardtii* cells exposed to 0.2 mM Ni was inhibited in the first 24 h of the treatment, compared to the control (Fig. 1, inset). In contrast, *C. reinhardtii* cells exposed to a concentration of 0.1 mM Ni showed similar growth to untreated cells during the 7-day experiment, with no significant differences. This data is in line with the study of Zheng et al., (2013), where the growth of *C. reinhardtii* exposed to Ni concentrations between 125-150 μM decreased considerably, whereas it showed similar growth to untreated cells at Ni concentrations between 30-90 μM. These results demonstrate once again the high tolerance of *C. merolae* to at least an order of magnitude higher concentrations of Ni compared to the microalgae present in neutral aquatic environments.

Cellular viability assessment by confocal microscopy allowed us to estimate the proportion of living cells present in the cell population exposed to 1-10 mM Ni (Fig. 2). In the case of 1 mM Ni, the number of viable cells remained nearly constant (∼90%) throughout the experiment, reaching a viability of ∼91% in day 15, which was somewhat lower than the proportion of viable control cells in day 15 (∼97%). From day 5, significant differences were observed between the untreated cells and cells exposed to Ni concentrations above 1 mM, and this difference was most noticeable in day 15 (p < 0.01), where the proportion of viable cells exposed to 6 mM and 10 mM Ni was a mere 4% and 2%, respectively. Interestingly, we observed the recovery of cell viability in the case of 3 mM Ni, i.e. from 56% in day 10 to 82% on day 15 (Fig. 2) confirming the long-term metabolic adaptability of *C. merolae* to such high Ni concentrations. The cell survivability data is in contrast with the work by Misumi et al., (2008) who observed up to 20% viability of *C. merolae* cells upon 10 mM Ni treatment on day 70, measured as autofluorescence of the chloroplast. In the latter work, the experimental and culture conditions were considerably different, as a different Ni source was used (Ni(NO_3_)_2_ instead of NiSO_4_); growth pH was slightly lower (2.3 compared to 2.5 in this work) and the final cell suspension volume was also different (50 versus 125 mL in this work), all of which may affect the effective toxicity of Ni in the cells. Moreover, in Misumi et al., (2008), the starting OD of the culture is not specified. In the present study, the starting OD_750_ was ∼0.150 at 750 nm, which could have caused much higher total effective toxicity of Ni compared to the work by Misumi and colleagues (2008).

**Figure 2.**
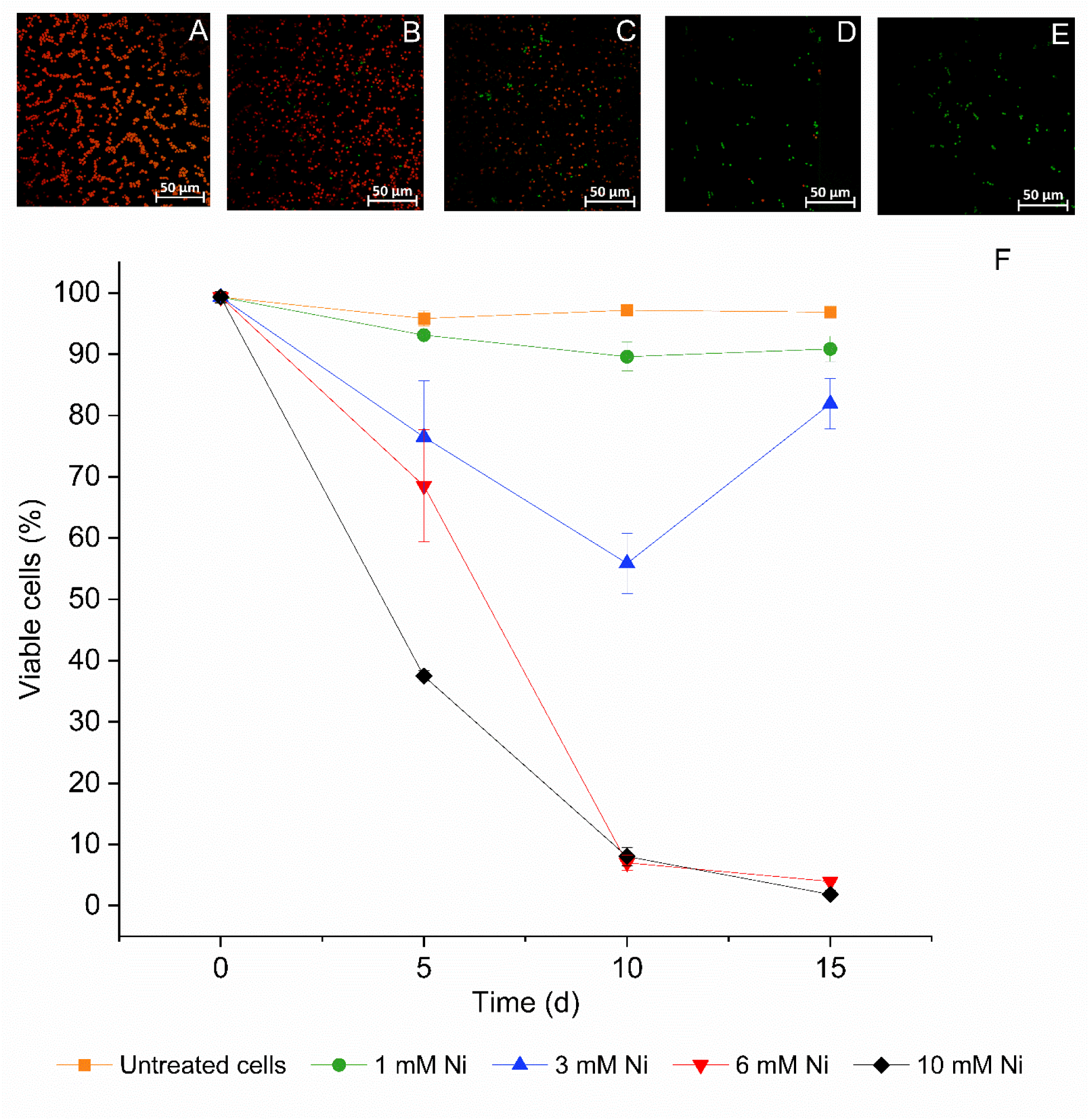
Confocal fluorescence imaging of viable *C. merolae* cells exposed to various nickel concentrations. Dual Laser measurements by CLSM were performed following the protocol by Millach et al. (2019). Viable (excitation at 555 nm and emission 585 nm) and non-viable (excitation at 405 nm and emission at 435 nm) cells are depicted in red and green, respectively. Cells were visualized on day 15 of the experiment. (A), untreated cells; (B), 1 mM Ni; (C), 3 mM Ni; (D), 6 mM Ni; and (E), 10 mM Ni. For each timepoint the viable cells were counted and the percentage of living cells in the whole cell population was calculated, and the corresponding viability data is shown in (F). Data in (F) represents the mean ± SD of two independent biological replicas (n=2). Scale bar: 50 μm.

### 2.2. Functional resilience of the C. merolae photosynthetic apparatus upon high nickel exposure

#### 2.2.1. Quantitative changes to photosynthetic and photoprotective pigments

The estimation of the total content of Chl *a* and carotenoids (Car) in the *C. merolae* cells exposed to different concentrations of Ni confirmed significant changes in the amount of both types of pigments during Ni exposure. After 15 days of up to 3 mM Ni exposure, the pigment content per cell was similar to the control sample (Fig. 3 and Tab. S1). However, at 6 mM Ni, the total Chl *a* and Car content decreased by 92% and 78%, respectively, compared to the untreated control, while at 10 mM Ni the values were beyond the detection threshold (Fig. 3 and Tab. S1). These observations are in line with the cell viability data (Fig. 2) which showed a significant decrease in the proportion of viable cells at 6 mM and 10 mM Ni. In higher plants, Ni treatment leads to dissociation of the entire photosynthetic complexes from the membranes (Szalontai et al., 1999) or their specific subunits. e.g., 17- and 24 kDa polypeptides stabilizing the oxygen evolving complex of PSII (Boisvert et al., 2007). In consequence, degradation of PS is likely triggered at the highest Ni concentrations applied in this study leading to the decrease of the total photosynthetic and photoprotective pigment content (Fig. 3). This is in contrast to other extremophilic microalgae, such as halophilic green alga *Dunaliella salina* in which total Car increase was observed during adaptation to Fe (Mojaat et al., 2008).

**Figure 3.**
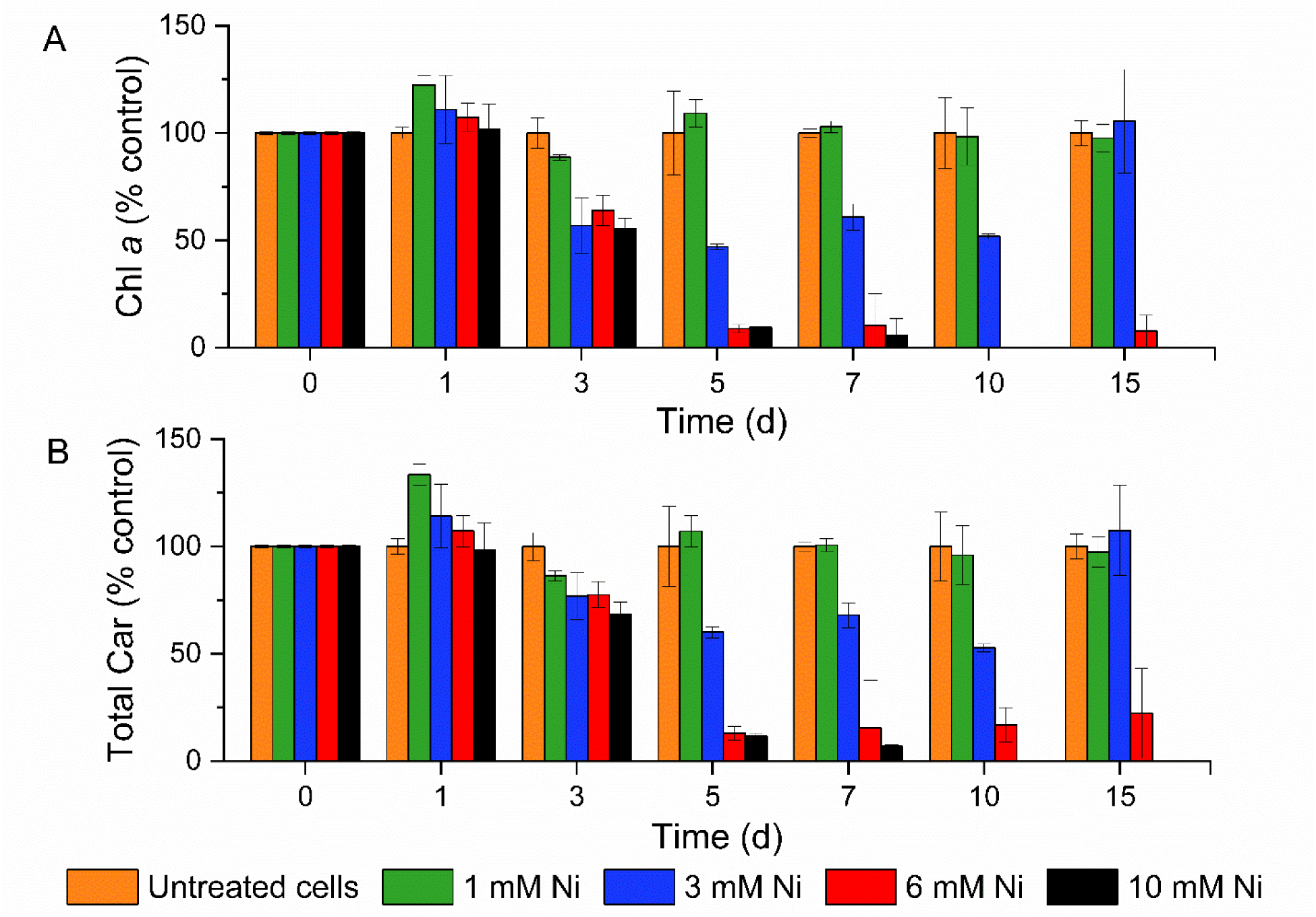
Quantification of total chlorophyll *a* (A) and carotenoid (B) content under cell exposure to different concentrations of nickel. The pigments were spectrometrically quantified in pg of pigment per cell, and are normalized to the total pigment content of the control (% control, Chl *a* 0.0250 – 0.0485 pg cell^-1^ and total carotenoids 0.077 – 0.0141 pg cell^-1^). Data represents the mean ± SD of two independent biological replicas (n=2) with a significant p-value < 0.05.

**Table 1.** PBS/PS ratio of untreated C. merolae cells and cells exposed to different Ni concentrations during 15 days of metal exposure. Data corresponds to the mean ± SD of two biological replicas (n=2).

| PBS/PS ratio | Untreated cells | 1 mM Ni | 3 mM Ni | 6 mM Ni | 10 mM Ni |
| --- | --- | --- | --- | --- | --- |
| DAY 0 | 1.09 ± 0.01 | | | | |
| DAY 5 | 1.07 ± 0.02 | 1.06 ± 0.01 | 1.14 ± 0.02 | 1.17 ± 0.000 | 1.16 ± 0.004 |
| DAY 10 | 1.15 ± 0.01 | 1.11 ± 0.001 | 1.06 ± 0.01 | 1.50 ± 0.019 | 1.33 ± 0.030 |
| DAY 15 | 1.18 ± 0.01 | 1.13 ± 0.002 | 1.13 ± 0.005 | 1.46 ± 0.041 | 1.10 ± 0.140 |

The effect of Ni on the decrease of photosynthetic and photoprotective pigments is well documented when this heavy metal is used in much lower (µM) concentrations than the Ni concentration range used in this study. For instance, the green microalga *Ankistrodesmus falcatus* decreased the total Chl *a* and Car content after 96 h treatment with 0.015-0.064 µM Ni (Martínez-Ruiz and Martínez-Jerónimo, 2015).Similarly, three different *Spirulina* species exhibited a severe decrease in Chl content after exposure to 10-100 µM Ni (Balaji et al., 2014). Therefore, our pigment quantification data demonstrates the remarkable integrity of *C. merolae* photosynthetic machinery and photosynthetic membranes during adaptation to up to 3 mM Ni, in contrast to other mesophilic and extremophilic phototrophs.

#### 2.2.2. Spectroscopic analysis of C. merolae under nickel exposure

The cell viability data upon exposure up to 3 mM Ni is in line with the OD_750_ data, where a cell growth recovery was observed. The same dynamics of cell growth recovery was observed by the measurement of room temperature (RT) absorption spectra of cell suspensions (Fig. 4). In contrast to the resilience of *C. merolae* to the Ni treatment, growth of a mesophilic green alga *C. vulgaris* was inhibited by 90% upon 15-45 μM Ni exposure (Santos et al., 2019). Moreover, the growth rate of *Desmodesmus sp.* was reduced by 18% when 0.19 mM Ni was used (Rugnini et al., 2017). On the other hand, in the case of the green microalga *Ankistrodesmus falcatus*, 2 μM Ni caused a total inhibition of microalgal cell growth (Martínez-Ruiz and Martínez-Jerónimo, 2015).

**Figure 4.**
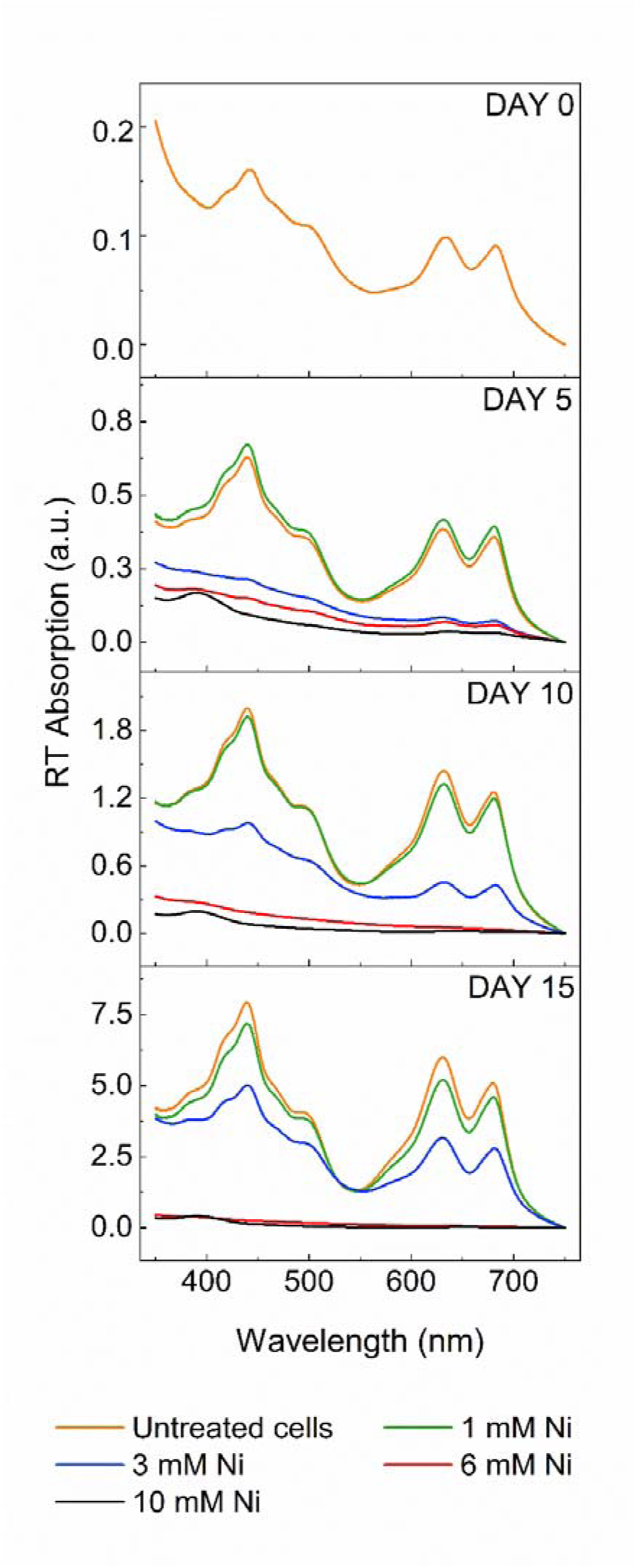
RT absorption spectroscopy of *C. merolae* cells upon long-term nickel adaptation. Spectra (averaged from 2 biological replicas) were obtained on days 5, 10 and 15 of the 1-10 mM Ni treatment.

The 77K Chl fluorescence analysis of the whole cells showed the relative changes in the functional antenna size of PSI (peaking at 726 nm) during long-term *C. merolae* adaptation to 1-10 mM Ni (Fig. 5A). On day 5, a 35-56% decrease in the antenna size of PSI was observed for the cells treated with 3, 6- and 10-mM Ni compared to the control and 1 mM Ni-treated cells. However, for 3 mM Ni-treated cells, the size of the PSI antenna recovered to the control level on days 10 and 15 of Ni treatment. Similarly, PSI fluorescence recovery was observed for 6 mM Ni-treated cells (Fig. 5A). In the case of 10 mM Ni treatment, no PSII or PSI fluorescence was detected from day 5 onward until the end of the experiment. Upon excitation of the PBS (at 600 nm), an increase of the 660 nm peak was observed on day 10, for 1- and 3-mM Ni-treated cells, and on day 15 for 3 mM Ni (Fig. 5B and Tab. 1), indicating the decoupling of PBS from PSII upon Ni treatment. The PSII peak was significantly blue-shifted in the case of the cells treated with 3-10 mM Ni (Tab. 2), indicating the occurrence of conformational changes (Boussac et al., 2020) and/or subunit or pigment dissociation from this complex (Andrizhiyevskaya et al., 2005) at such high Ni concentrations. A similar phenomenon was observed for the spinach PSII complex subjected to thermal stress (Wang et al., 2017).

**Figure 5.**
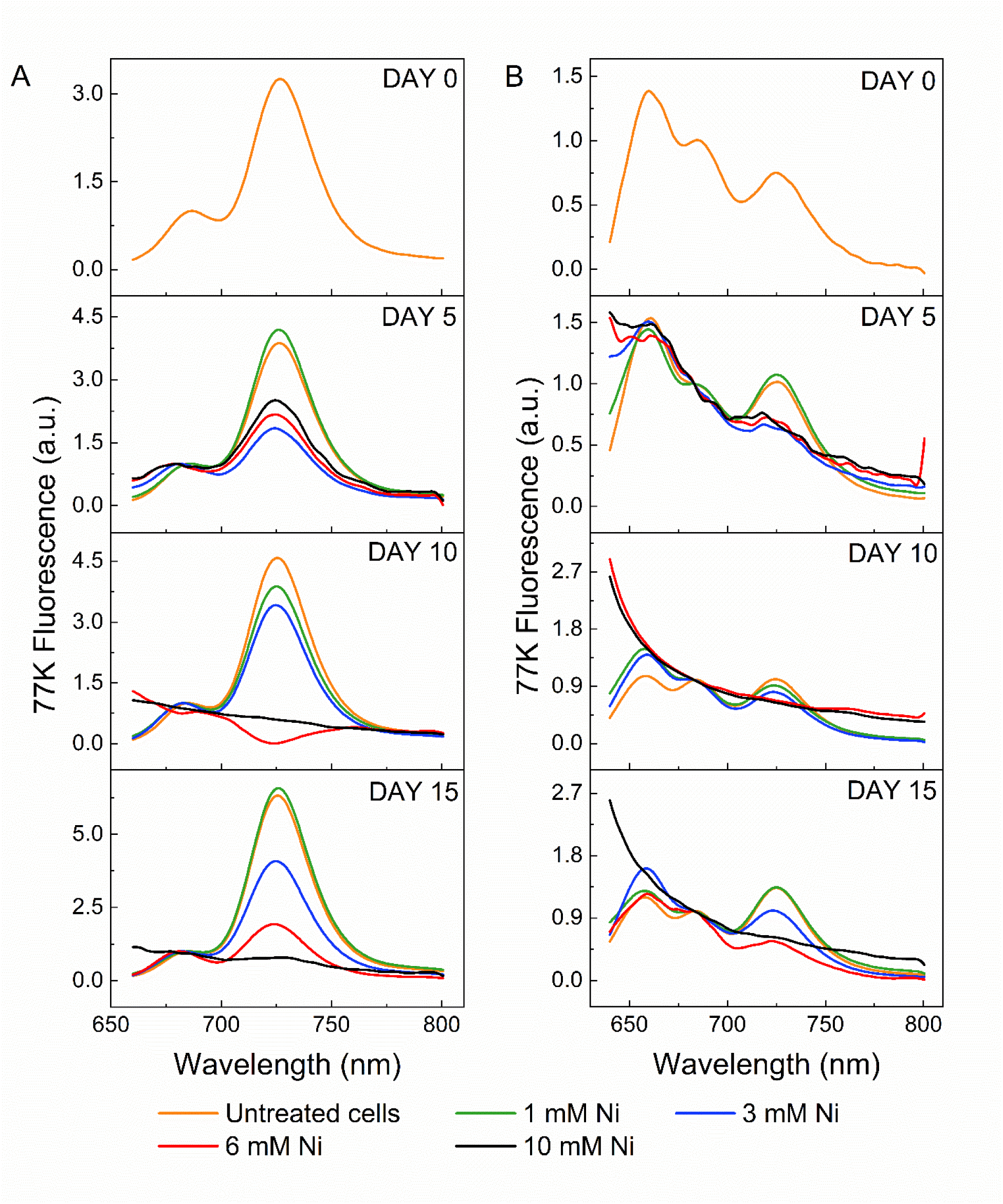
77K fluorescence spectroscopy of the *C. merolae* cells long-term adapted to various Ni concentrations. (A) corresponds to the emission spectra using Chl *a* excitation at 435 nm on days 5, 10 and 15, respectively. (B) corresponds to the emission spectra upon PBS excitation at 600 nm on days 5, 10 and 15, respectively. All spectra were normalized to the PSII peak (685 nm) and were averaged from 2 biological replicas.

**Table 2.** 77K Fluorescence maxima of PSII in untreated cells and cells exposed to various Ni concentrations. Data corresponds to the mean ± SD of two biological replicas (n=2).

| Time (day) | Untreated cells | 1 mM Ni | 3 mM Ni | 6 mM Ni | 10 mM Ni |
| --- | --- | --- | --- | --- | --- |
| 5 | 686 | 686 | 681* | 680* | 680* |
| 10 | 685 | 683 | 683 | 680.5* | - |
| 15 | 686 | 685 | 682.5* | 681* | - |

\*Significant PSII fluorescence maximum blue shift ( $p < 0.05$ ).

#### 2.2.3. Photosynthetic performance of C. merolae under nickel stress

The presence of heavy metals including Ni inside the photosynthetic cells can directly affect the efficiency of the photosynthetic electron transfer at many levels including structural destabilization and catalytic inhibition of the donor side of PSII, i.e., by replacing Ca in the Mn_4_O_5_Ca cluster and extrinsic PsbO subunit (Yocum, 2008), replacing Mg in the Chl molecules, replacing heme and non-heme Fe or replacing Cl in the PSII complex structure (reviewed in Nowicka, 2022). Other effects of heavy metal treatment include aberration of the thylakoid membrane organization (e.g. membrane de-stacking), membrane lipid peroxidation leading to the formation of ROS and oxidative damage of lipids, proteins and other biomolecules and direct inhibition of Calvin-Benson-Bassham cycle enzymes including RuBisCo and phosphoribulokinase (reviewed in Shahzad et al., 2018).

Of all the multi-level deleterious effects of heavy metals, it is important to assess the performance of the photosynthetic apparatus under this type of stress, especially in the context of the capability of the primary energy production during long-term adaptation to metal stress. One of the main and most important parameters that determine photosynthetic performance is the maximal PSII quantum yield of PSII (F_v_/F_m_ parameter), which corresponds to maximum efficiency at which light absorbed by PSII is used for photochemistry (reduction of strongly bound quinone Q_A_) (Baker, 2008). Several studies of the Ni effect on PSII performance have been conducted in microalgae, albeit applying a much lower Ni concentration range (0.05-2 mM Ni) than those used in the present work due to the lethal effect of high concentrations of this heavy metal on model algal systems such as *Euglena gracilis* (Ahmed and Häder, 2010), *Phaeodactylum tricornutum* (Guo et al., 2022), *Chlamydomonas reinhardtii* (Zheng et al., 2013) and *Scenedesmus obliquus* (Mallick and Mohn, 2003).

As shown in Fig. 6A, F_v_/F_m_ values of 0.57, 0.57 and 0.52 were obtained for control, 1- and 3-mM Ni treated cells, respectively, confirming the existence of the efficient photochemical process in PSII during cellular adaptation up to 3 mM Ni. In contrast, at 6- and 10-mM Ni concentrations, an irreversible decline of the F_v_/F_m_ parameter was observed already after 24 h Ni exposure. Consequently, the effective operating efficiency of PSII (Y(II) parameter, Fig. 6B) showed a similar trend, with values of 0.5 for the untreated, as well as 1- and 3-mM treated cells,0.46 and 0.42 respectively. This observation indicates that almost half of the absorbed quanta of light are effectively converted into photochemical energy in the PSII reaction centers during cellular adaptation to up to 3 mM Ni. As expected, the Y(II) parameter decreased to zero from day 10 of the 6- and 10-mM Ni treatment. The shut-down of the photosynthetic apparatus in the presence of 6- and 10-mM Ni (from day 10) is clearly demonstrated by the measurement of the Y(NO) (Fig. 6C) and Y(I) (Fig. 6D) parameters corresponding to non-regulated non-photochemical quenching (NPQ) (fraction of energy passively dissipated as heat and fluorescence) and effective quantum yield of PSI, respectively.

**Figure 6.**
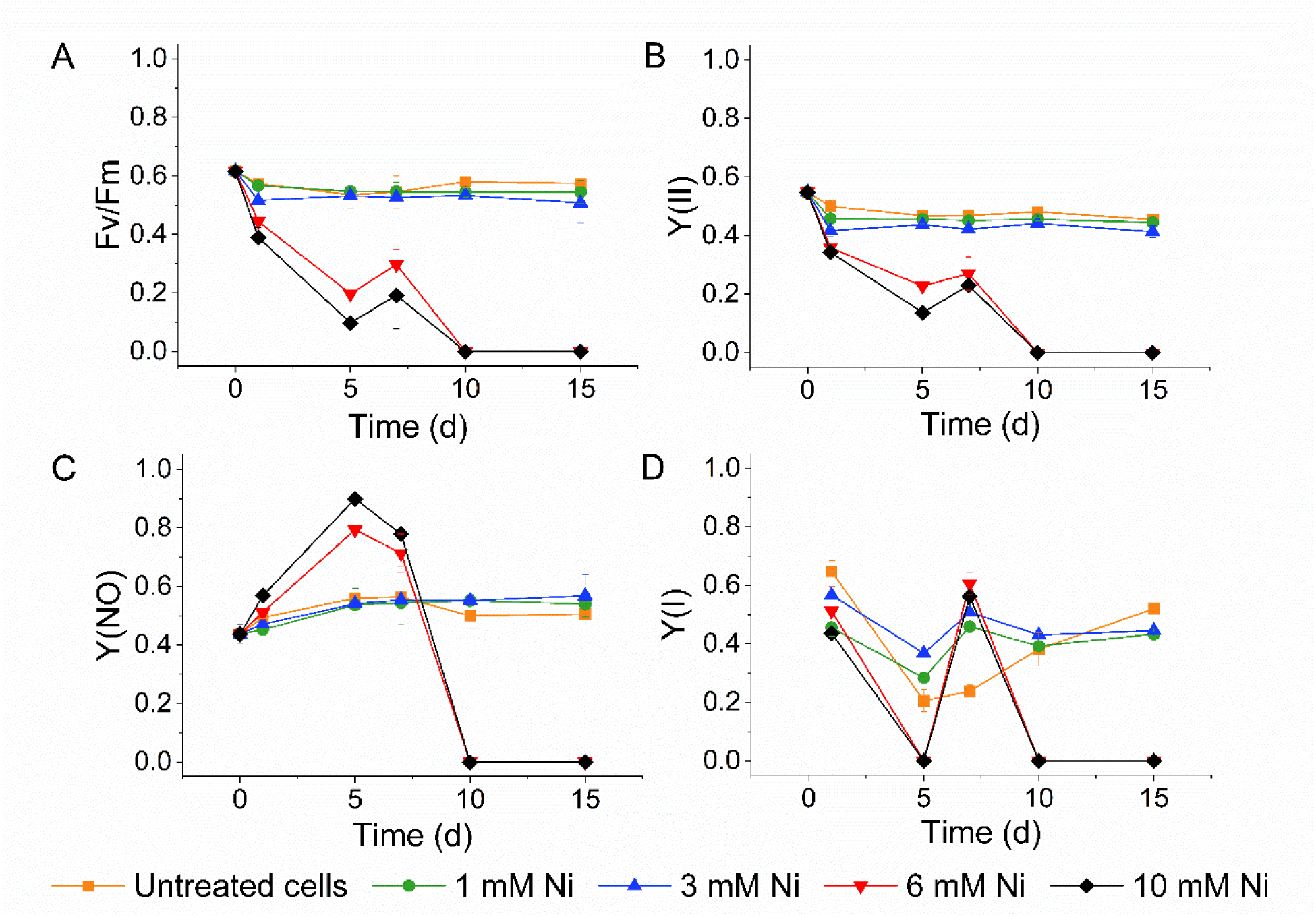
DUAL-PAM analysis of photosynthetic performance in *C. merolae* cells long-term adapted to various nickel concentrations. Shown are: maximum efficiency of PSII (F_v_/F_m_ ratio; A), effective photochemical efficiency of PSII (Y(II); B), non-regulated NPQ (Y(NO); C) and the effective photochemical efficiency of PSI (Y(I); D). Data for all parameters is the mean ± SD of two independent biological replicas (n=2).

We examined the efficiency of the regulated NPQ which is a crucial fast photoprotective process triggered in PSII upon acidification of luminal pH during exposure of photosynthetic cells to various types of long-term stress including high light (Horton and Hague, 1988; Adams and Demmig-Adams, 1992; Kirilovsky and Kerfeld, 2013; Krupnik et al., 2013; Ruban, 2016; Bassi and Dall’Osto, 2021) and heavy metals (Zhang et al., 2020; Nowicka, 2022). The *in vivo* investigation of the NPQ kinetics (Fig. 7) during long-term adaptation of *C. merolae* cells to up to 3 mM Ni displayed the kinetic curves that were similar to the previously observed for cyanobacterial counterparts exposed to high light stress (Calzadilla and Kirilovsky, 2020). The exposure of dark-adapted cells to low intensity of blue light triggered a transient decrease in the maximum fluorescence of PSII (F_m_ʹ), which corresponds to State 2-to-State 1 transition present in all the PBS-containing phototrophs including *C. merolae* (Canonico et al., 2020; Abram et al., 2022). The latter process constitutes a fast adaptive strategy that evolved in phototrophs to balance the photosynthetic electron flow between both photosystems under any stress conditions that may affect the redox state of the plastoquinone pool (Kargul and Barber, 2008). This observation is in line with the dynamic remodeling of the functional antenna size of PSI observed in this study (Fig. 5). Therefore, our study for the first time confirms the existence of efficient photoprotective responses, i.e. regulated NPQ and state transitions, in the *C. merolae* cells upon their long-term exposure to high (up to 3 mM) concentration of Ni.

**Figure 7.**
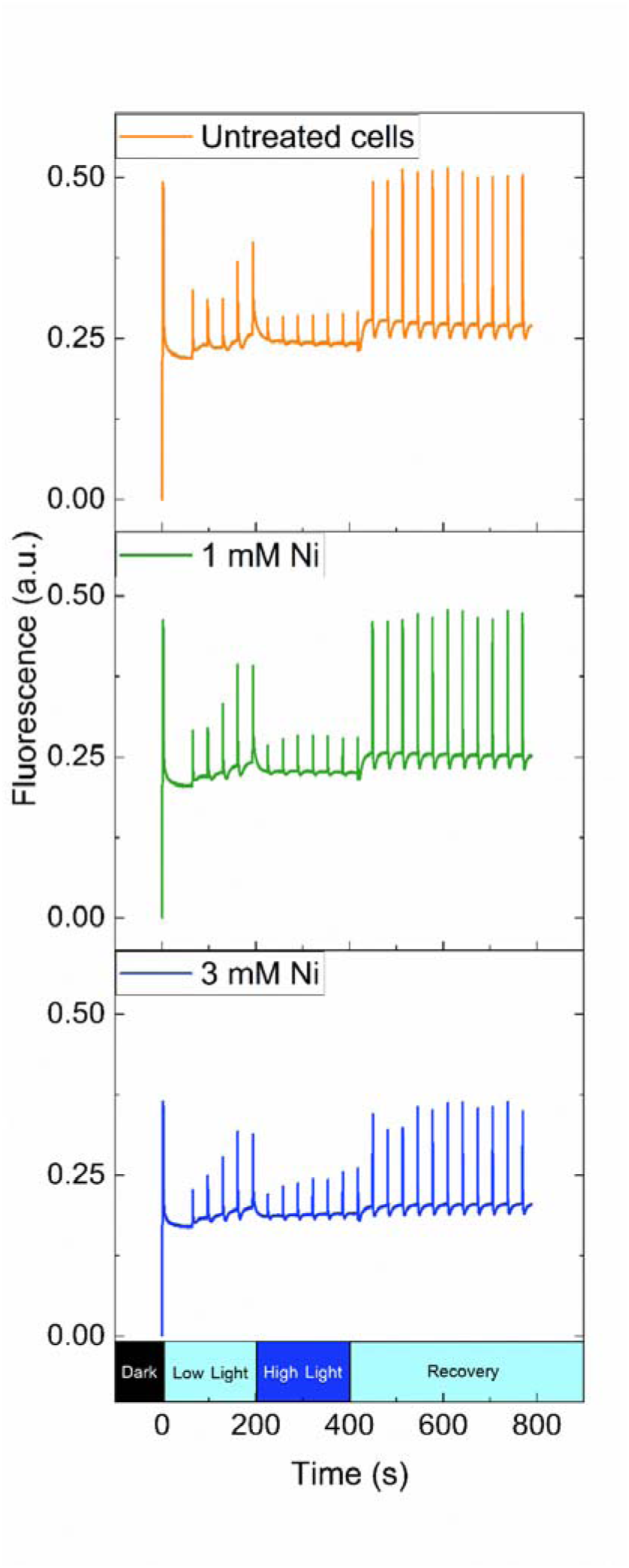
Fluorescence induction kinetics analysis of the *C. merolae* cells exposed to various nickel concentrations. After a long-term acclimation process (15 days), untreated cells, 1 mM Ni and 3 mM Ni treated cells were first dark adapted and then exposed to a sequence of different light pulses, as described in Materials and Methods. The ‘Low Light’ bar corresponds to a period of 200 s of low intensity blue light illumination (90 µE m^−2^ s^−1^); ‘High Light’ bar corresponds to a period of 200 s of high intensity blue light illumination (1500 µE m^−2^ s^−1^); while ‘Recovery’ bar corresponds to a 400 s period of low intensity blue light illumination (90 µE m^−2^ s^−1^).

#### 2.2.4. Analysis of oxygen evolution and consumption in C. merolae cells upon Ni exposure

To further assess the integrity of the photosynthetic apparatus of *C. merolae* during long-term adaptation to high concentrations of Ni we performed the analysis of the oxygen evolution and consumption activities in cell suspensions exposed to 1-10 mM Ni during 15 days. The overall oxygen evolution rate (measure of PSII photochemical activity) in *C. merolae* cells exposed to 6- and 10-mM Ni was significantly higher (461% and 509% of the control) compared to the untreated samples (Fig. 8A) up to 7 days of Ni exposure followed by a decrease of this parameter on days 10-15 (133% and 227% of the control). This observation is in line with the PAM data (Fig. 4). The molecular basis underlying the transient increase of O_2_ production during adaptation to high Ni concentration is presently unknown. However, it is possible that the inhibition of the competing reactions, such as water-water cycle, light-induced mitochondrial respiration and/or photorespiration may be responsible for this phenomenon (Raven et al., 2020). This intriguing hypothesis will be investigated in the near future.

**Figure 8.**
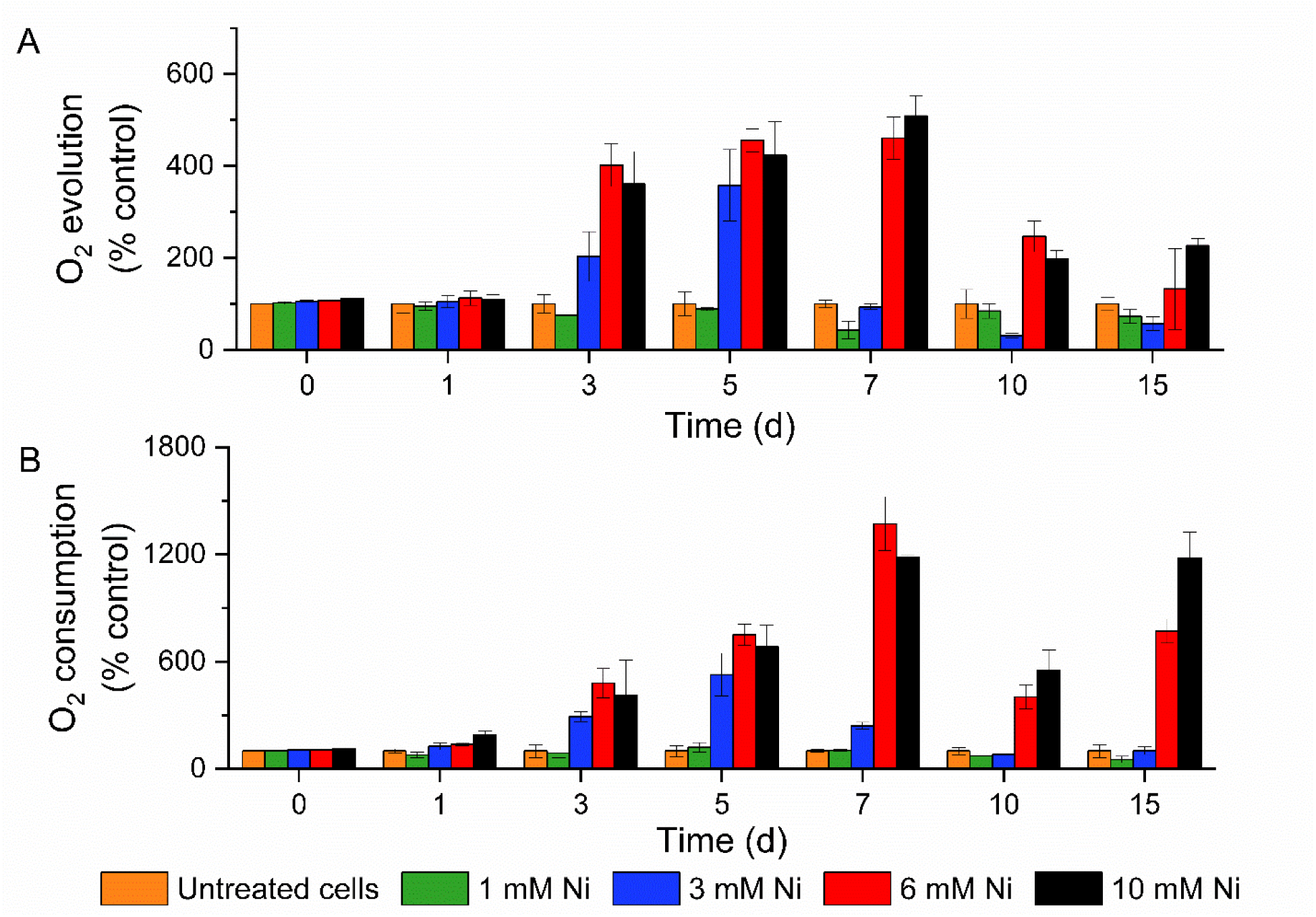
Oxygen evolution (A) and basal consumption (B) of *C. merolae* cells during long-term adaptation to various nickel concentrations. Values are expressed as nmoles of O_2_ per 10^6^ cells per hour, and normalized to the control (% control, O_2_ evolution 0.78 – 2.07 nmol O_2_ 10^6^ cells^-1^ h^-1^ and O_2_ consumption 0.21 – 2.18 nmol O_2_ 10^6^ cells^-1^ h^-1^). Data is the mean ± SD of two independent biological replicas (n=2).

We observed the highest O_2_ consumption rate (measure of mitochondrial respiration) on day 7 in the presence of 6 mM Ni (Fig. 8B). We hypothesized that the increase of this parameter could be due to impairment or remodeling of the mitochondrial respiratory chain components upon Ni treatment. To shed light on this issue, we applied 2,4-dinitrophenol (DNP) to the *C. merolae* cells adapted for 7 days to 3- and 6-mM Ni. DNP is a well-known mitochondrial protonophore that dissipates pH gradient across the mitochondrial membrane leading to uncoupling of oxidative phosphorylation from the mitochondrial electron transport chain (Goldgof et al., 2014).We assumed that if the respiratory chain components underwent the putative functional or structural impairment due to the intracellular Ni action, the DNP treatment would not further increase their activity and thus, the overall oxygen consumption. However, the DNP treatment of Ni-adapted cells resulted in markedly higher oxygen consumption rates in 3 and especially in 6 mM Ni-adapted cells (1.4- and 7.7-fold increase for 3- and 6-mM Ni samples, respectively; Fig. S2) most likely due to remodeling hypothesized before and increase in the number of respiratory chain components. We propose that the increased amount of the respiratory chain molecular components serves as the efficient adaptive strategy to maintain the cellular energy homeostasis during long-term adaptation of *C. merolae* to Ni stress. The verification of this intriguing hypothesis will be addressed in the near future.

#### 2.2.5. ROS production in C. merolae cells long-term exposed to nickel

Metal intoxication increases ROS levels as they play a significant role in many ROS-producing mechanisms, including the Haber–Weiss cycle, Fenton’s reactions, and disruption of the photosynthetic electron chain leading to superoxide and singlet oxygen production and reduction of the glutathione pool (Pinto et al., 2003). ROS also act as signaling molecules that have an impact on a network of antioxidants, level of antioxidant enzymes, and other stress related molecules (Shcolnick and Keren, 2006). The well-established effects of heavy metals on cellular metabolism include oxidative damage of proteins, lipids and DNA due to the generation of superoxide (O_2_^•-^), hydroxyl radicals (OH^•-^), singlet oxygen (^1^O_2_) and hydrogen peroxide (H_2_O_2_) which at the cellular level evoke oxidative stress by disturbing the redox homeostasis in plant and algal cells (Mallick and Mohn, 2000; Hossain et al., 2012; Sytar et al., 2013; Rezayian et al., 2019). In consequence, protein and lipid degradation occurs together with ion leakage, oxidative DNA damage, redox imbalance, and degradation of the cellular membranes, ultimately leading to programmed cell death (Nagajyoti et al., 2010).

The role of Ni in generation of ROS-induced oxidative stress in algal cells is well documented (reviewed in Nowicka 2022). We therefore examined the Ni-induced adaptability of *C. merolae* to ameliorate the cellular ROS effects by measuring the rates of ROS production at different timepoints of Ni adaptation and expressing those relative to the ROS production rate in the untreated cells (Fig. 9). We observed the significantly highest increase of ROS production rate on day 5 for all Ni treatments (p < 0.01). This data confirms that during long-term adaptation of *C. merolae* to high Ni concentration, the ROS level is inherently high, even in the first days of cellular adaptation to 1- and 3-mM Ni when the cell growth and viability are relatively high compared to the control (Fig. 1 and 2). Despite the generally higher levels of ROS in Ni-adapted cells compared to the non-stressed cells, the ROS amelioration metabolic machinery is capable of reducing their level, as shown by the sinusoidal character of the ROS production curves. We propose that the ROS amelioration pathways are likely to be highly active in *C. merolae* cells throughout the high Ni adaptation process. Verification of this intriguing possibility is currently underway.

**Figure 9.**
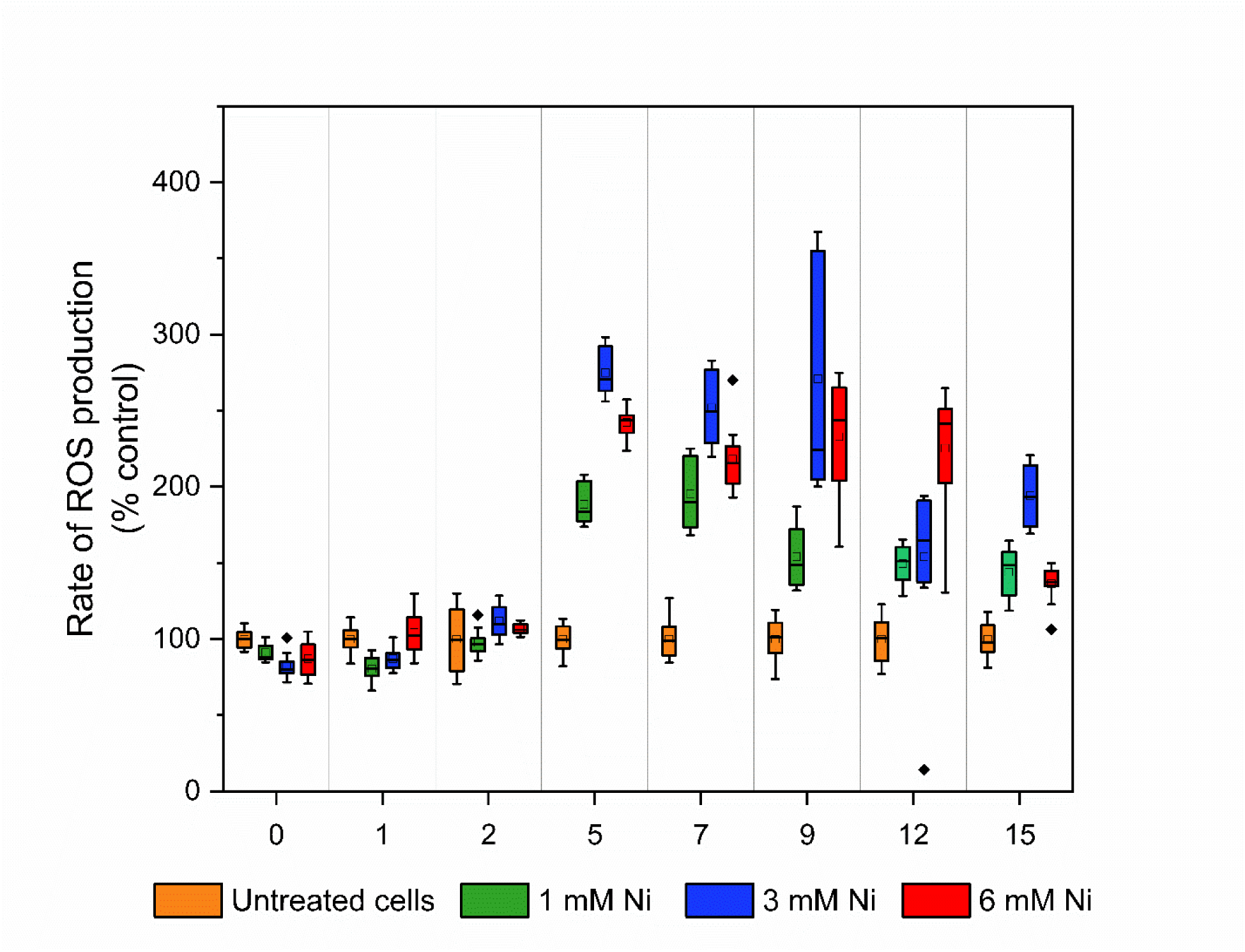
Analysis of ROS production in *C. merolae* cells during long-term exposure to various nickel concentrations. The rate of ROS production was measured for each sample using the fluorescent probe, 2’,7’-dichlorodihydrofluorescein diacetate. Values correspond to the rates of ROS production normalized to protein concentration (µg mL^-1^) and then expressed as a percentage of Ni untreated control (% control, 12.51 – 43.51 rate of ROS production per protein (µg mL^-1^)). The colored boxes represent the interval data between 25-75% of the total values. The upper and lower whiskers correspond to the maximum and minimum values excluding the outliers. The black horizontal lines inside the boxes are the medians, while the squares are the average values and the rhombi correspond to the outlier values. Data is the mean ± SD of two independent biological replicas, with 6 technical replicas for each biological replica (n=2).

#### 2.2.6. Association of nickel with C. merolae cells

At the cellular level, the defense mechanisms to a metal ion can be either exclusion of the metal from the cell, the uptake and modification of the metal to a less toxic form, followed by the metal’s extrusion from the cell, or by internal sequestration (Hall and Williams, 2003). These processes are a result of a coordinated network of biochemical processes, which increase the cell’s ability to maintain metabolic homeostasis, and minimize oxidative stress. A common strategy for heavy metal detoxification is metal compartmentalization and sequestration in different cellular organelles such as vacuoles or as electron dense bodies in the cytosol, chloroplasts and mitochondria (Sharma et al., 2016). In various microalgal species, the vacuole is the main locus for sequestration of metals. Thus, in *Pseudochlorococcum typicum* the exposure of cells to Pb caused the formation of spherical electron-dense bodies in the vacuole (Shanab et al., 2012). On the other hand, *C. reinhardtii* and *E. gracilis* microalgae sequester Cd complexes in the chloroplast and, in the case of *E. gracilis* additionally in the mitochondrion (Mendoza-Cózatl and Moreno-Sánchez, 2005; Hanikenne et al., 2009).

The absence of typical vacuoles in *C. merolae* combined with its remarkable adaptability to high concentrations of Ni (as shown by this study) implies that this alga may have evolved other efficient mechanisms of detoxification of this heavy metal. Interestingly, *C. caldarium,* another member of Cyanidiales that contains large vacuoles, has been shown to deposit high levels of metals such as Fe, Al and Zn in cytoplasmic electron dense bodies (Nagasaka et al., 2004). In the case of *C. merolae*, no data has been reported to date on the exact localization of Ni within the cellular compartments or its putative association with the cell membrane during long-term adaptation to this heavy metal. To gain insight into this issue, a combined electron microscopy and elemental analysis was performed on *C. merolae* cells long-term exposed to various concentrations of Ni at different timepoints using three independent approaches. The combination of these three analyses at different time points suggests that the main strategy of *C. merolae* is to actively extrude Ni from the cells, as the majority of this heavy metal was associated with the cell membrane (Fig. 10, 11 and Tab. 3).

**Figure 10.**
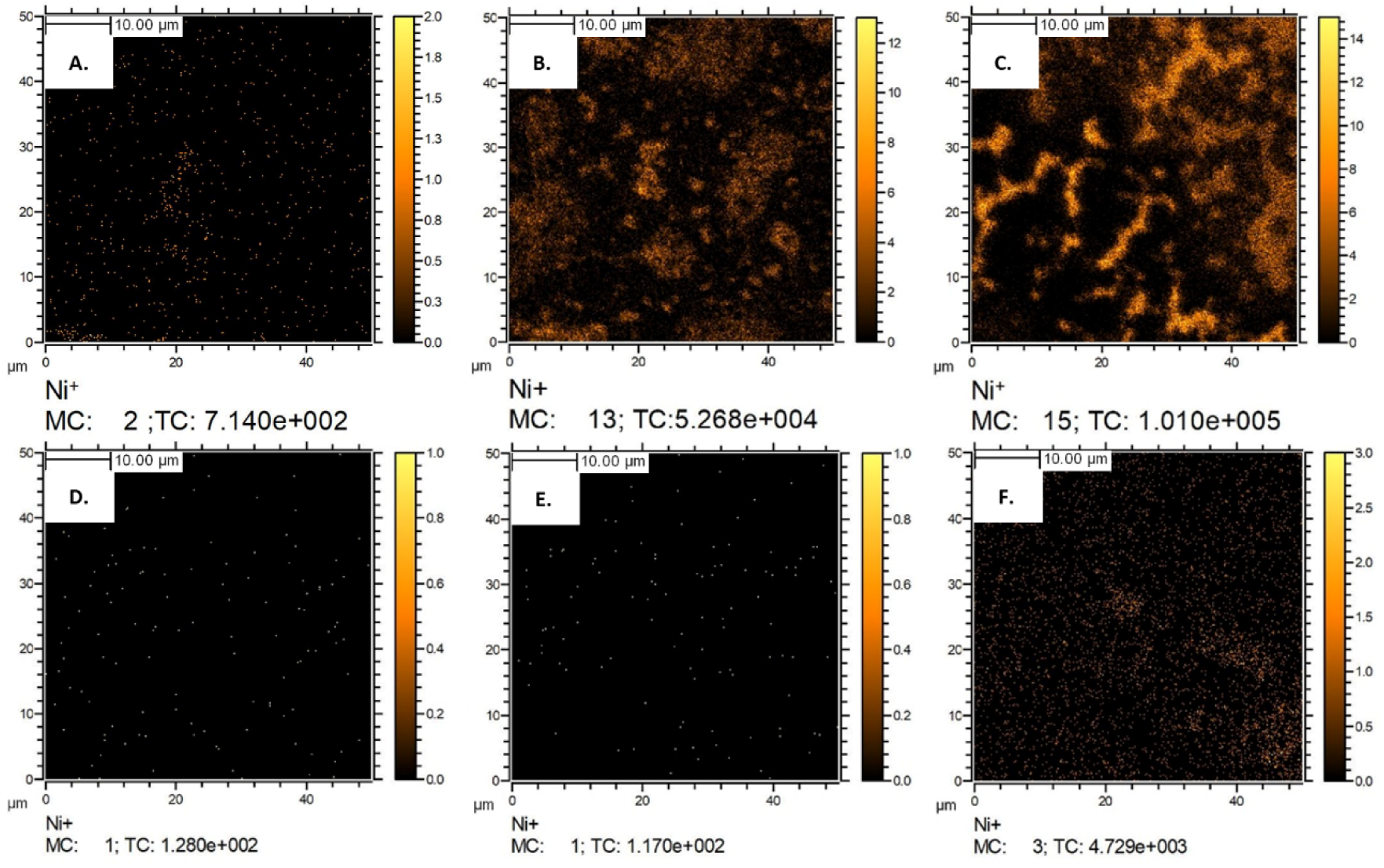
Nickel ion maps of the *C. merolae* cells during long-term adaptation to Ni. Micrographs correspond to a 10 µm × 10 µm area of immobilized *C. merolae* cells exposed to varying concentration of Ni. A-C, Ni maps of the samples without any processing; D-F, Ni maps of the samples subjected to the one-step washing. The concentration of Ni^2+^ in the growth medium was 1 mM (A, D), 3 mM (B, E), and 6 mM (C, F). The color scale bar on the right of each figure shows the number of Ni ion counts collected at the corresponding point. Scale bar: 10 µm. MC – maximum number of counts per pixel (also shown as a scale), TC - total number of counts. The small white spots correspond to random spike noise signal.

**Figure 11.**
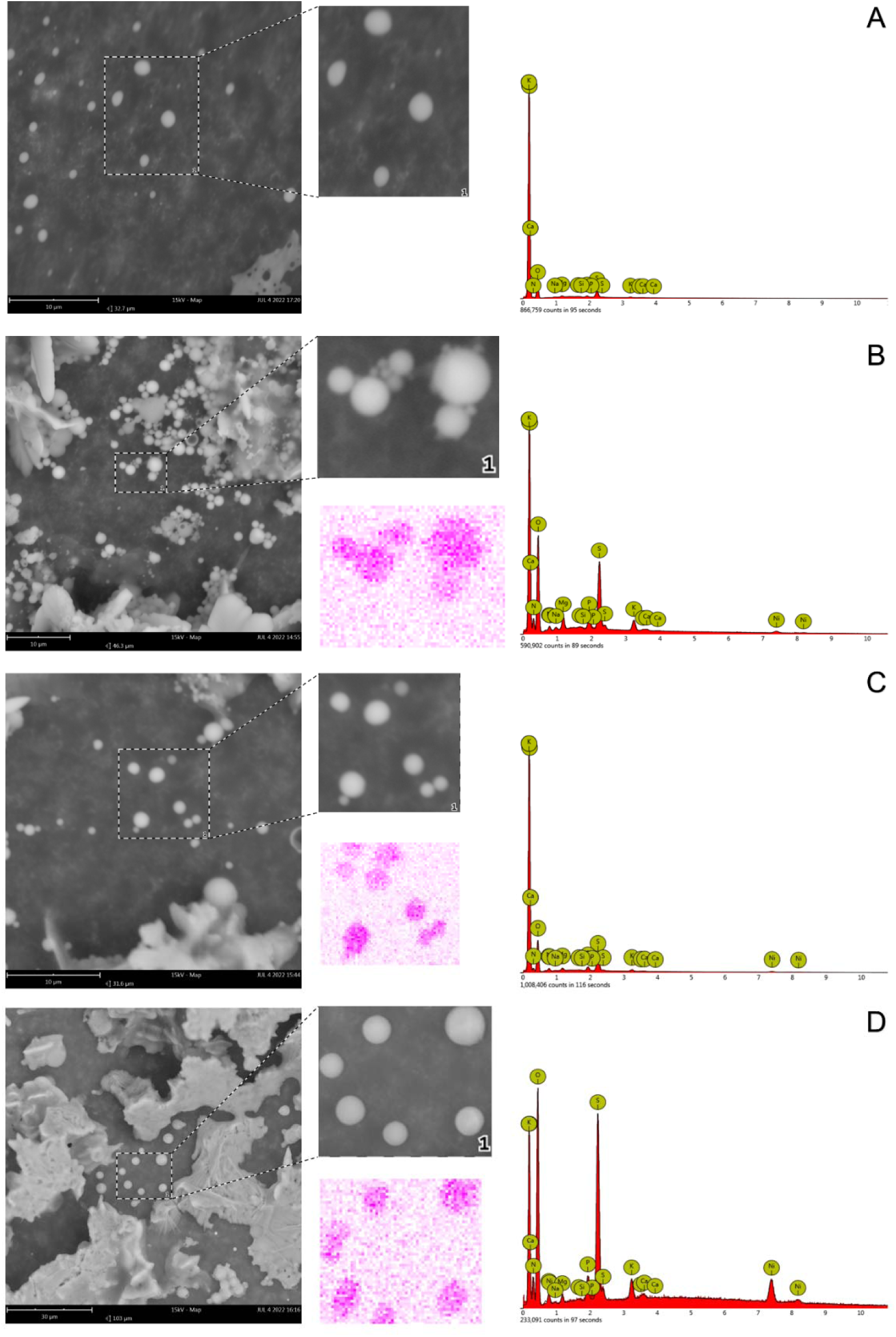
SEM-EDS imaging of Ni in *C. merolae* cells long-term adapted to various nickel concentrations. Each panel corresponds to a different treatment (day 5 of Ni exposure), either (A) untreated cells; (B) 1 mM Ni; (C) 3 mM Ni; and (D) 6 mM Ni treated cells. For each panel shown is: micrograph with the scale bar (left), the magnified spot of selected cells to be analyzed, the EDS micrograph of Ni (center); and the spectra of the EDS map (right). Scale bar: 10 μm (A, B, C) and 30 μm in (D).

**Table 3.**
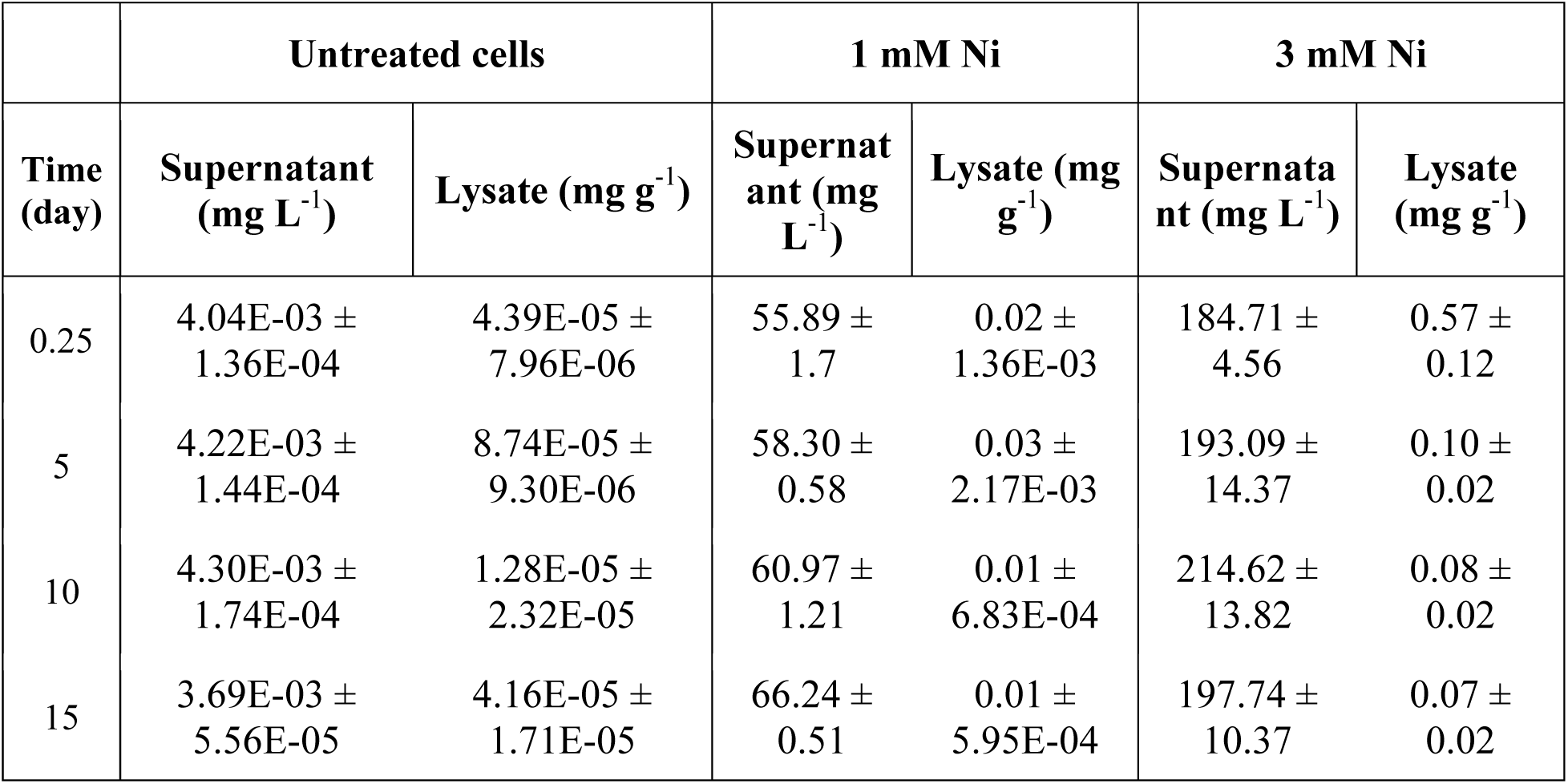
Nickel quantification by ICP-MS in *C. merolae* cells long-term adapted to various Ni concentrations. Quantification of Ni is shown for the growth medium (supernatant; mg Ni L^-1^) and the lysate (mg Ni g^-1^ of wet biomass). Data corresponds to the mean ± SD of two independent biological replicas (n=2).

The elemental analysis based on Inductively Coupled Plasma Mass Spectrometry (ICP-MS) showed that for all the Ni concentrations applied less than 1% of the total Ni added to the growth medium is associated with the cells (Tab. 3). As expected, the amount of Ni present in the cell lysate at 3 mM Ni is more than 3-fold higher compared to the samples exposed to 1 mM Ni for all the time points except for day 5 samples (2.8-fold increase, Tab. 3). The level of Ni present in the lysate of 1 mM Ni treated cells is constantly low at different timepoints (0.01-0.03 mg g^-1^), demonstrating that the cells were able to efficiently adapt to this concentration of metal throughout the treatment despite the cyclic increase of the cellular ROS level (Fig. 9). We hypothesize that the low level of cellular Ni is due to basal active extrusion of this heavy metal or other mechanisms that prevent the Ni accumulation inside the cells and its deleterious effect on cell metabolism. This hypothesis is supported by the physiological data showing no significant differences in the growth curves or photosynthetic parameters at 1 mM Ni compared to the untreated cells (Fig. 1 and 6). In contrast, the Ni content in the cell lysate at 3 mM Ni decreases progressively during 15-day treatment from 0.57 in the first 6 hours to 0.10 mg/g and 0.07 mg/g on days 5 and 15 of the treatment, *respectively* (Tab. 3). We postulate as metabolic fine-tuning occurred during the first 5 days of the treatment that promoted inducible Ni extrusion and additional Ni defense mechanisms (over and above the basal Ni active extrusion) at such high Ni levels in the growth medium. This hypothesis is in line with the physiological data of partial inhibition of cell growth at 3 mM Ni within the same timeframe followed by the recovery phase (Fig. 1). In the case of 6- and 10-mM Ni, due to severe inhibition in the cell growth (Fig. 1) and cell viability (Fig. 2) the elemental analysis on these samples was not possible.

In an independent approach based on Time-of-Flight Secondary Ion Mass Spectrometry (TOF-SIMS) we confirmed that the majority of Ni is loosely associated with the *C. merolae* cell membrane. SIMS is a destructive method of analysis. Due to the interactions with the environment (matrix effect) and the higher probability of emission, the former metal ions are predominantly the single charged particles (Benninghoven, 1994). Therefore, the obtained Ni^+^ distribution maps can be equated with occurrence the signal from the Ni signal associated either with *C. merolae* cells and/or with Ni deposited during evaporation of growth medium.

Fig. 10 presents the Ni^+^ ion emission SIMS-TOF maps registered for the *C. merolae* cells immobilized on the silicon substrate following their 15-day exposure to 1-6 mM Ni. The maps presented in the upper row (Fig. 10A-C) correspond to the samples without any washing processing (see Materials and Methods). Upon increasing Ni^2+^ concentration in the bulk of solution the increase in the amount of Ni deposited on the cells is clearly visible. In contrast, no significant Ni signal was observed for the cells subjected to the washing procedure (Fig. 10D-F). In the latter case, the signal is much weaker indicating that most of Ni visible in Fig. 10A-C was physically adsorbed on the cell surface. The low intensity of the observed signal may be explained by nature of the SIMS-TOF technique. The sampling depth of this technique is approximately 1-3 nm (Muramoto et al., 2012),thus the ion maps represent the signal collected from the cell surface.

The SIMS spectra were analyzed for the signals from the substrate (Si^+^) and from the algal cells (Fig. S3). The organic fragment, C_3_H_8_N^+^, was chosen as representative SIMS marker for the organic matter of the cells as it corresponds to the product of peptide ionization (Schönwälder et al., 2014; Thiruvallur Eachambadi et al., 2021)and its mass is close to the mass of the main isotope of Ni. The Ni content for low-concentration (1 and 3 mM Ni) cultures correlates with the presence of the organic ion (Fig. S4) indicating the adsorption of this ion on the cell surface. However, for the 6 mM Ni-treated samples some differences were observed. Spots of the local Ni accumulation, which do not correlate with the presence of organic structures, were detected (Fig. S3). This may indicate that a significant portion of Ni remains in the solution in the ionic form.

The association of Ni with the cell membrane was additionally confirmed by the analysis based on Scanning Electron Microscopy combined with Energy Dispersive X-Ray Spectroscopy (SEM-EDS), whereby the majority of Ni is detected on the cell surface (Fig. 11).

#### 2.2.7. Structural changes to thylakoid organization during C. merolae adaptation to high nickel concentration

Heavy metal exposure may affect substantial changes to thylakoid membrane organization as shown for the red alga *C. caldarium* exposed to Al (Nagasaka et al., 2002) or higher plants exposed to Cd (Parmar et al., 2013). These changes are due to alteration of the membrane integrity through lipid peroxidation under metal stress, degradation of protein components embedded within the thylakoids and alteration of the ionic milieu affecting the dynamics of membrane stacking and unstacking. *C. merolae* has a simple cellular ultrastructure, with up to ⅔ of the cell volume occupied by a single chloroplast under non-stress conditions. In this microalga, thylakoids do not form grana or stromal lamellae but are organized into a network of loosely positioned membranes (Toyoshima et al., 2016; Miyagishima and Tanaka, 2021).

To observe the putative changes of the thylakoid architecture during long-term adaptation of *C. merolae* do Ni stressor we performed Transmission Electron Microscopy (TEM) analysis of the cellular ultrastructure on day 7 of 1-6 mM Ni treatment (Fig. 12). We observed no significant differences in the number of thylakoids or their spatial organization in the 1 mM Ni samples compared to the untreated control (Fig. 12A/D and 12B/E). However, during adaptation of *C. merolae* cells to Ni concentrations above 1 mM, structural changes were observed in chloroplasts. In 3 mM Ni-treated cells, the number of thylakoids was significantly lower compared to the control and 1 mM Ni-treated cells (Fig. 12C, F and G), despite high cell viability (Fig. 2) and high photosynthetic activity (Fig. 6) at this Ni concentration. At 6 mM Ni, it was not possible to calculate the number of thylakoids due to the strong aberration of the chloroplast structure. Not only the number of thylakoids layers was affected by exposure to 3- and 6-mM Ni but also the chloroplast size: the cells adapted to 1- and 3-mM Ni showed a slightly shorter longitudinal axis compared to the control sample (2.32 *versus* 2.14 and 2.15 µm for untreated cells and cells exposed to 1- and 3-mM Ni, respectively) after 15 days of metal exposure (p < 0.01), as estimated by confocal fluorescence imaging. In the case of 6- and 10-mM Ni it was not possible to estimate the chloroplast size after 15 days of experiment due to the low viability and the fact that most of the cells are disrupted.

**Figure 12.**
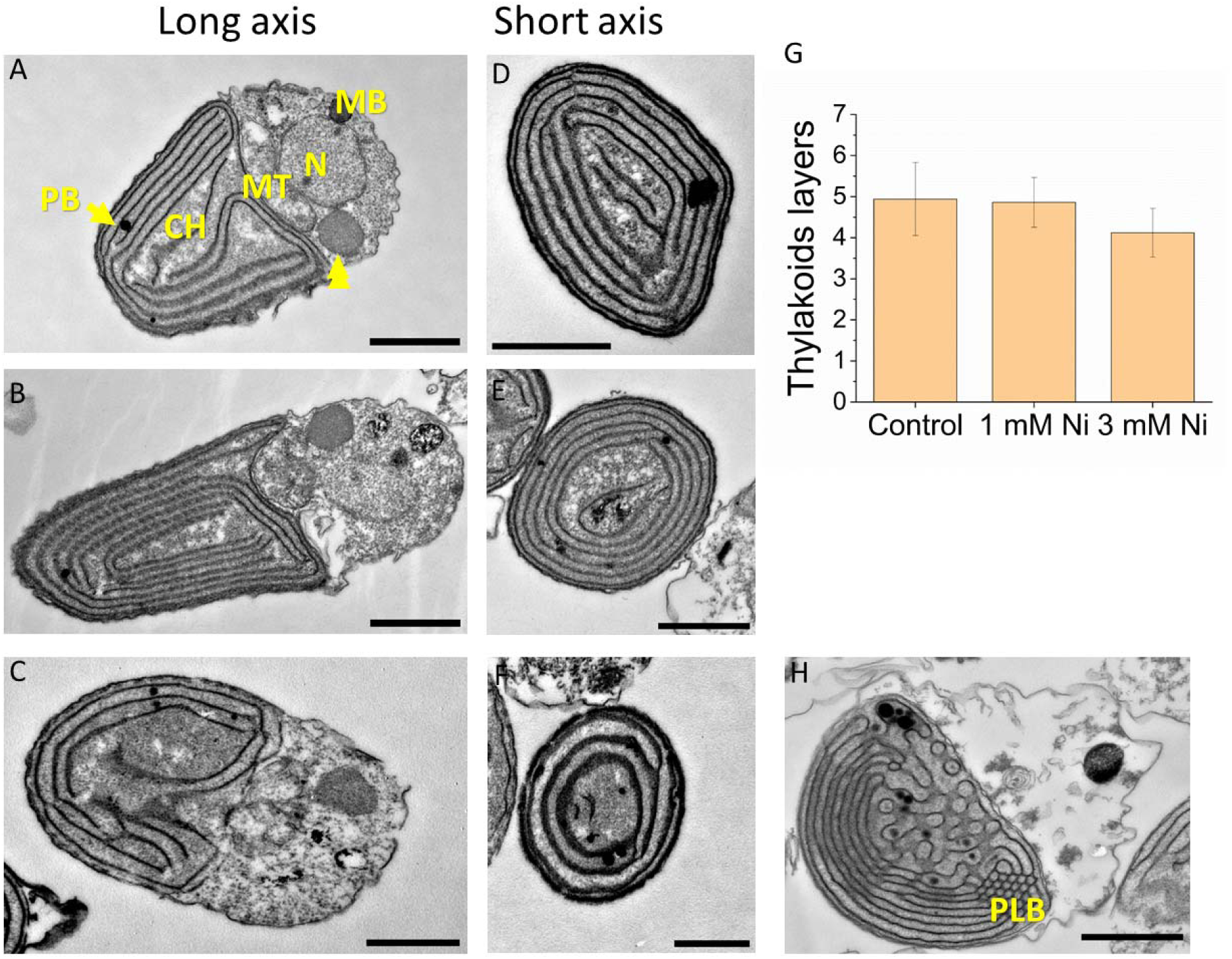
TEM imaging of *C. merolae* cells during long-term adaptation to various nickel concentrations. (A, D), untreated cells; (B, E), cells exposed to 1 mM Ni; (C, F), cells exposed to 3 mM Ni; and (H), cells exposed to 6 mM Ni for 7 days. For each Ni treatment, cells are visualized along the long (A, B, C) and short axes (D, E, F). (G) Quantification of the number of thylakoids on day 7 of the experiment. Data corresponds to the mean ± SD following the analysis of 50 cells (per each sample) using two independent biological replicas (n=2). N: nucleus, CH: chloroplast, MT: mitochondrion, PG: plastoglobule, PLB: prolamellar body, MB: microbody, arrowhead depicts a vacuole-like organelle. Scale bar: 1 µM.

The characteristic feature of the chloroplasts exposed to 6 mM Ni was the presence of tubular prolamellar body (PLB)-like structures (Fig. 12H). The PLBs constitute the lipid reservoirs for developing thylakoids (Bykowski et al., 2020) and have been shown to prevent lipid peroxidation and RNA damage under oxidative stress conditions (Almsherqi et al., 2009; Solymosi and Mysliwa-Kurdziel, 2021). The PLBs have been observed in association with both cyanobacterial thylakoids and eukaryotic chloroplasts. They represent a unique non-lamellar, but cubic-phase inner membrane structures. In spite of the similar lipid composition, major differences are observed in the protein composition of chloroplast thylakoids and PLBs (reviewed Solymosi and Mysliwa-Kurdziel 2021) with the latter containing much higher lipid:protein ratio and high content of the light-dependent NADPH:protochlorophyllide oxidoreductase (LPOR), and enzyme catalyzing protochlorophyllide photoreduction. The lipid-dependent formation of the photoactive LPOR oligomers was proposed to be a mechanism inducing the formation of the PLBs (Gabruk et al., 2017; Fujii et al., 2019). We postulate that the formation of PLBs in the *C. merolae* chloroplasts subjected to 6 mM Ni stress may constitute the efficient adaptive mechanism triggered in response to high Ni stress to sustain efficient Chl biosynthesis, which is required for the photosynthetic components repair.

Another structural feature observed in this study for Ni-adapted chloroplasts is the increased number of plastoglobules (PGs) which are lipoprotein structures participating in chloroplast metabolism and stress responses (Arzac et al., 2022). On average, the number of PGs in the chloroplast’s cells adapted to 1- and 3-mM Ni increased around 2-fold compared to the control after 7 days of Ni exposure, whereas in the chloroplasts adapted to 6 mM Ni the increase was 6-fold compared to the control (Fig. 12). For 10 mM Ni it was not possible to count the number of PGs due to the low cell viability impaired intactness at such high metal concentration.

PGs form spherical lipoprotein particles that are associated with thylakoid membrane zones of high curvature (Austin et al., 2006). The characterization of PG proteome, composed of ∼ 30 core proteins with multiple functions, revealed an unexpectedly active metabolic role of these structures in contrast to the previously postulated role of PG as the passive lipid bodies of the storage function (Vidi et al., 2006; Ytterberg et al., 2006). Overall, PG metabolome and proteome components are related to four major metabolic pathways, i.e., Chl degradation, thylakoid remodeling, biosynthesis of prenylquinones as well as Car biosynthesis and metabolism (Lundquist et al., 2012; van Wijk and Kessler, 2017; Michel et al., 2021; Arzac et al., 2022). It is well documented that the quantity and size of PGs increase in response to many types of biotic and abiotic stresses (reviewed in Venzhik et al., 2019). These include heavy metal toxicity (Sandalio et al., 2001; El-Banna et al., 2019), high light (Lichtenthaler et al., 1981), temperature stress (Zhang et al., 2010), leaf senescence (Hörtensteiner, 2006), high salinity (Naidoo et al., 2011), desiccation (Pressel and Duckett, 2010; Fernández-Marín et al., 2013) and pathogens (Raman et al., 2006). Such active involvement of PGs in various stress adaptation processes is underpinned by their diverse functional and metabolic roles. Therefore, the increase of the number of PGs observed in this study is in line with the active metabolic role of these stress-marker organelles during *C. merolae* adaptation to high concentrations of Ni.

## Conclusions

In this work, we have performed comprehensive analysis of the long-term adaptive responses to extremely high Ni stress (in mM range) in the cells of a model extremophilic (acido-thermophilic) microalga *C. merolae* isolated from the heavy metal-rich environments of volcanic origin. The complementary spectroscopic, microscopic and elemental analyses allowed us to dissect several molecular mechanisms underlying the long-term adaptation mechanisms to high Ni concentration. These include: (i) extrusion of Ni from the cells and lack of significant Ni accumulation inside the cells as demonstrated by the microscopic and elemental SIMS, SEM-EDS and ICP-MS analyses; (ii) maintaining the efficient photoprotective responses including NPQ and state transitions (up to 3 mM Ni) to ensure the highest photosynthetic performance, as shown by DUAL-PAM and 77K fluorescence analyses, and limit, by cutting down light absorption, the generation of ROS and other redox-active molecules that are released despite these photoprotective mechanisms; (iii) dynamic remodeling of the chloroplast ultrastructure such as formation of metabolically active PLBs and PGs together with loosening of the thylakoid membrane architecture as shown by TEM analysis; (iv) activation of ROS amelioration metabolic pathways as demonstrated by the dynamic regulation of ROS production and relatively high cell viability up to 3 mM Ni; and (v) maintenance of the efficient respiratory chain functionality likely by increasing the amount of the respiratory chain components as demonstrated by the marked stimulation of oxygen consumption in response to pharmacological uncoupling of the mitochondrial membranes with DNP. All the dynamic processes identified in our study underlie the capability of *C. merolae* to adapt efficiently to extremely high Ni levels that exceed by several orders of magnitude the levels of this heavy metal in the natural environment of the microalga. By dissecting the molecular mechanisms of the remarkable adaptability of *C. merolae* to high Ni concentration, our study paves the way for the biotechnological application of this extremophile in heavy metal recycling and engineering new traits in higher plants and microalgae for high heavy metal tolerance/sequestration.

## Materials and Methods

### Cell culturing, cell growth measurement and IC_50_

*Cyanidioschyzon merolae* strain NIES-3377 (obtained from the Microbial Culture Collection of the National Institute for Environmental Studies in Japan) was cultivated in a modified 2x Allen medium (Allen, 1959) at 42°C, pH 2.5 and with shaking at 110 rpm (Minoda et al., 2004). Cell suspensions (125 mL) were grown in glass conical flasks (Duran®) in continuous white light of 90 µE m^-2^ s^-1^ using an FL40SS-ENW/37H growth chamber (Panasonic). The Ni stock solution was prepared by adding nickel sulphate hexahydrate (NiSO_4_ x 6H_2_O) in MilliQ water to obtain a final concentration of 200 g L^-1^. An appropriate volume of 2x Allen’s culture medium, *C. merolae* stock culture and Ni stock solution was added to each flask to obtain a final concentration of 1, 3, 6- and 10-mM Ni. Each volume was pre-calculated so as to maintain the final Ni concentration. In addition, a control culture was prepared using the same procedure but omitting the Ni solution. Ni adaptation experiment was carried out for a final of 15 days. Exponentially growing cells were employed for all experiments, with a starting OD_750_ ∼0.15. For all the timepoints and Ni concentrations investigated in all the subsequent analyses, the samples were analyzed in parallel during the same measurements using two independent biological replicas (n=2). Cell growth was monitored daily by recording room temperature (RT) absorption spectra in the range of 370-750 nm and recording the values of OD_750_ using a UV-1800 spectrophotometer (Shimadzu).

The IC_50_ is the Ni concentration after 15 days at which the growth of *C. merolae* is inhibited by 50 % as compared to untreated cells. The growth inhibition as percentage was calculated on the OD_750_ values after 15 days of growth of the two-biological replica for each treatment (Monteiro et al., 2011).

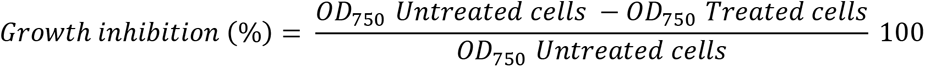

The IC_50_ value was calculated using OriginPro software (OriginLab Corporation, 2018) through nonlinear curve Dose/Response.

### Confocal fluorescence imaging

An LSM700 confocal laser scanning microscope (Zeiss) was used to acquire the confocal images of the cells at 63x magnification. Prior to imaging, cells were immobilized on microscopic slides using a 0.75 mm layer of 2% agar that was cut into squares. An aliquot (10 µL) of cell suspension was placed over the agar square, then covered with a coverslip. The average size of chloroplasts (measured along the longest axis) was estimated within at least 50 cells for two independent biological replicas. For cellular imaging, two different channels were simultaneously used to visualize PSI and PSII (488 nm excitation wavelength and 516 nm fluorescence emission) as well as PBS complexes (555 nm excitation wavelength and 585 nm fluorescence emission). To determine the cell viability (the number of viable and non-viable cells a dual laser beam setup was used, as described in (Millach et al., 2019). Firstly, a red channel (555 nm excitation wavelength and 573 nm fluorescence emission) was used to observe viable cells and the chloroplast autofluorescence signal. At the same time, a green channel (405 nm excitation wavelength and 435 nm fluorescence emission) was applied to visualize non-viable cells and the non-photosynthetic autofluorescence signal. Four different images were obtained for every biological replica. At least 80 cells for each replica were evaluated for cell viability. Cell counting and statistical analyses were performed to assess the exact number of viable and non-viable cells in the culture exposed to each Ni concentration.

### Pigment quantification

For the total Car and Chl *a* quantification, pigments were extracted from the cells using neat (≥ 99.8 %) N,N-dimethylformamide (DMF). Briefly, an aliquot of cell suspension was first centrifuged for 10 min at 14,000 g. After the removal of the supernatant, 1 mL of neat (≥ 99.8 %) DMF was used to extract pigments from the cell pellet in the dark at -20°C for 24 h. The samples were mixed vigorously in the dark, then centrifuged for 10 min at 14,000 g. RT absorption spectra were measured in the range of 350-750 nm using a UV-1800 spectrophotometer (Shimadzu) using pure DMF as blank. The following equations (1) and (2), with extinction coefficient for Chl *a* and Car in DMF of 83.9 L g^-1^ cm^-1^ (Moran, 1982) and 2500 L g^-1^ cm^-1^ (Hirschberg and Chamovitz, 1994) respectively, were used to calculate the concentration of pigments (μg mL^-1^) in the cellular extract taking into consideration the dilution factor (Dil) and the ratio between the volume of the sample (V_sample_, mL) and the volume of DMF (V_DMF_, mL) used to extract the pigments (Wellburn, 1994; Borella et al., 2022):

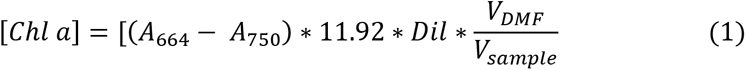

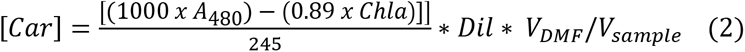

where A is the absorbance at an indicated wavelength. The pigment content (pg mL^-1^) was normalized to the number of cells per mL, as determined with a hemocytometer (Burker), and expressed as % control.

### RT Absorption spectroscopy

RT absorption spectra were acquired in a range of 350-750 nm with a 10 mm-optical path length using a UV-1800 spectrophotometer (Shimadzu) supplied with a TCC-100 temperature-controlled cell holder.

### 77K Fluorescence spectroscopy

Steady-state fluorescence emission spectra of intact cells were measured at 77K using an LS55 Fluorescence Spectrometer (Perkin Elmer), as described by Krupnik et al. (2013). Emission spectra of Chl *a* and phycobilisomes (PBS) were recorded using excitation wavelengths of 435 and 600 nm, respectively. The emission spectra were normalized to the PSII peak (685 nm) Each spectrum is obtained from the average of two independent biological replicas.

### ROS production measurement

To measure the rates of ROS production inside the cells, 2’,7’-dichlorodihydrofluorescein diacetate (CM-H2DCFDA) was used, dissolved in DMSO to obtain a final concentration of 5 mM. In parallel, an aliquot (3 mL) of *C. merolae* cell suspension was centrifuged ed for 8 min at 1,000 g and the supernatant was removed. The pellet was then resuspended in 3.5 mL of phosphate buffered saline (PBS: sodium chloride, 145 mM (0.85%) in phosphate buffer, 150 mM) (pH 7.0) at 30°C. An aliquot (6.5 µl) of DCF stock was added to the final concentration of 9.3 µM and the sample was gently vortexed. The cell suspensions (0.25 mL) were then transferred to a 48-well microplate. Oxidized probe fluorescence was measured with an Infinite M200 plate reader (TECAN, Männedorf, Switzerland) in real time for up to 65 min. The quantitative fluorescence data was processed with a Magellan^TM^ software (TECAN). Total protein content was measured with a Bradford method (Bradford, 1976). For each sample, 0.2 mL of cell suspension was mixed with 0.05 mL of 100 mM NaOH, 0.3 mL of the Bradford reagent and 1 mL of water in a 1.5 ml cuvette. Absorbance at 595 nm was measured using freshly prepared calibration curves and bovine serum albumin as a standard. Protein content (mg mL^-1^) was calculated considering a 1-cm optical path length and the extinction coefficient of 6.6 (g/100 mL)^-1^ cm^-1^. The ROS production rate was normalized to the protein amount in each sample using 2 biological replicas and 6 technical replicas for each biological replica.

### Oxygen evolution and consumption measurement

A Clark-type electrode (Hansatech) was used to determine the oxygen evolution and consumption rates in cell suspensions. Measurements were performed at 30°C using a freshly calibrated electrode. The sample (1 mL) was preincubated in the dark with gently stirring for 25 min after which oxygen consumption rate (nmol O_2_ h^-1^) was measured. The sample was then illuminated for 15 min at 2,000 µE m^-2^ s^-1^ using a KL 2500 LED light box (Schott) to measure the oxygen evolution rate (nmol O_2_ h^-1^). The rate values were then normalized on 10^6^ cells per mL, as determined by counting cells using a hemocytometer (Burker), and expressed as % control. For the oxygen consumption measurements in presence of 23.1 μM 2,4 dinitrophenol (DNP), 1 mL sample was dark adapted for 15 min after which the oxygen consumption rate (nmol O_2_ h^-1^) was measured. To determine the effective concentration of DNP for uncoupling the oxidative phosphorylation from the mitochondrial electron transport chain 100 μL aliquots of 0.1 mM DNP stock solution was added to cell suspensions in the dark every 10 min and the oxygen consumption rate was measured immediately (nmol O_2_ h^-1^). The rate values were normalized to 10^6^ cells per mL and expressed as % control.

### DUAL-PAM measurement

To measure the photosynthetic performance parameters, a dual-wavelength pulse-amplitude-modulated fluorescence (DUAL-PAM) was recorded using a DUAL-PAM 100 fluorimeter (Heinz Walz Gmbh, Germany) and a DUAL-PAM v1.19 software. An aliquot (1 mL) of cell suspension was placed in a 10 mm quartz glass cuvette supplemented with a magnetic stirrer and incubated for 10 min in the dark. Then, an actinic pulse (12 µE m^-2^ s^-1^) was applied to record the minimal fluorescence (F_o_: the fluorescence yield of the open reaction centers of PSII, after dark adaptation). The maximum fluorescence yield of PSII (F_m_) was then recorded by applying a saturation pulse (SP) of 4,000 µE m^-2^ s^-1^ for 300 ms at 620 nm. Maximum quantum yield of PSII (F_v_/F_m_), was calculated as follows:

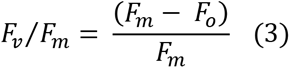

To quantify P700^+^ a dual wavelength unit (830/875 nm) was used to record simultaneously the changes in absorption at 700 nm (Klughammer and Schreiber, 1994). P_m_, defined as the maximal P700 change, was detected after application of a far-red light at 720 nm for 10 s, followed by a SP. For recording the Slow Kinetics Curves, a blue actinic light pulse of 90 µE m^-2^ s^-1^ was used followed by SPs applied every 30 s until a steady state was achieved. For each SP, the values of F_m_’ and P_m_’ (maximum PSII fluorescence yield and maximum change of the P700 signal in a given light, respectively; see Eq. 5-7), was recorded. Then, the actinic light (Blue Light; λ = 460 nm) was switched off. Effective quantum yields of PSII (Y(II)) and PSI (Y(I)) and non-regulated energy dissipation yield (Y(NO)) were calculated using the following equations:

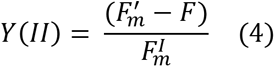

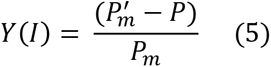

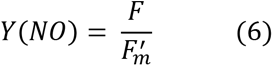

where Y(II), and Y(NO) are the effective quantum yield of PSII and the quantum yield of non-regulated energy dissipation, respectively, while Y(I) corresponds to the effective quantum yield of PSI. Besides, *F* is the fluorescence emission; *P* is the signal from the PSI reaction center before the application of the saturation pulse.

### Measurement of NPQ kinetics

The non-photochemical quenching (NPQ) was measured using a DUAL-PAM 100 fluorometer (Heinz Walz GmbH, Germany) by applying a standard procedure used for cyanobacteria, as described in (Canonico et al. 2021). Cells were dark adapted for 10 min with gentle stirring prior to the measurements. The F_m_ values of dark-adapted samples were registered by applying a saturating pulse (4,000 µE µE m^−2^ s^−1^, 400 ms, λ = 620 nm). The cells after 120 s of dark adaptation were subjected to a low intensity blue light illumination (90 µE photons m^−2^ s^−1^, λ = 460 nm). After 120 s the blue light intensity was increased to 1,500 µE m^−2^ s^−1^, λ = 460 nm) for 200 s. A recovery phase was also recorded whereby the actinic light intensity was decreased (90 µE m^−2^ s^−1^, λ = 460 nm). The maximal value of fluorescence was estimated by a multiple turnover flash (red light λ = 620 nm, 4,000 µE m^−2^ s^−1^, duration 400 ms) every 30 s.

### Inductively Coupled Plasma Mass Spectrometry (ICP-MS) analysis

Aliquots of cell suspensions (50 mL) were collected at different timepoints (6 h, and 5, 10 and 15 days) and centrifuged for 20 min at 2,500 g at 4°C. The supernatant (growth medium) was separated from the cell pellet and immediately processed by ICP-MS analysis. Alternatively, cell pellets were resuspended in a 2 mL aliquot of 0.05% SDS and 0.05 M 1,4-dithiothreitol, then boiled for 5 min and immediately used for ICP-MS analysis. An inductively coupled plasma mass spectrometer (NexION 300D ICP Mass Spectrometer, Perkin Elmer SCIEX, USA) was used. A conventional Mainhardt nebulizer and a quartz cyclonic spray chamber were used for sample introduction. The ICP-MS conditions were presented in Table 4. Nickel content was determined by ICP-MS monitoring the ^61^Ni isotope. The background interferences from the plasma gases, air entrainment and solvent were corrected by subtraction of reagent blank signals.

**Table 4.** Operating conditions for ICP Mass Spectrometer.

| Parameters | Value |
| --- | --- |
| Nebulizer Gas Flow | 0.83 L/min |
| Auxiliary Gas Flow | 1.15 L/min |
| Plasma Gas Flow | 16.00 L/min |
| ICP RF Power | 1350 W |
| Spray chamber | quartz type Scott |
| Sample cone | Nickel |
| Skimmer cone | Nickel |

For sample dilution and preparation of standards were used ultrapure water (MilliQ, Millipore) and nitric acid (HNO_3_, Sigma Aldrich). The reagent blank solution contained 1% of concentrated HNO_3_. Single standard solution containing Ni was prepared in reagent blank solutions. External calibration was performed with the standard solutions containing 0.1 µg/L, 1 µg/L, 10 µg/L, 100 µg/L of nickel. The limit of detection for nickel was 0.1 µg/L.

### TOF-SIMS

Time-of-Flight Secondary Ion Mass Spectrometry (TOF-SIMS) analysis was performed using a TOF-SIMS 5 spectrometer (IONTOF GmBH, Germany). To obtain good horizontal resolution and at the same time high spectral resolution, delayed extraction mode was used. The bismuth ion source was used operating at conditions: Bi_3_^+^, 30 keV, ion current: 0.08-0.12 pA over 50 × 50 μm^2^ area rastered at a resolution of 256 × 256 pixels. Mass spectra were monitored up to the m/z value of 900 and were internally calibrated using Na^+^, Si^+^, C_3_H_4_^+^, C_3_H_5_^+^, C_4_H_7_^+^ and C_5_H_9_^+^ ions. Two protocols of sample preparation for SIMS-TOF measurements were followed: (*i*) sample with cells placed in 2-Allen and Ni^2+^ solution after placing on silicon wafer was evaporated and then placed in SIMS-TOF chamber, or (*ii*) sample, before placing on silicon wafer, was washed in a single step using the following procedure: cells were centrifuged for 5 min at 268 x g at 21°C, then resuspended in 2-Allen medium without Ni^2+^ ions. A 5 µl volume of the cell suspension was subsequently placed on the silicon surface. After solvent evaporation the samples were immediately analyzed.

### Scanning Electron Microscopy with Energy Dispersive Spectroscopy analysis

Cells were collected and gently mixed with 8% dimethyl sulfoxide (DMSO). A 10 µL aliquot of cell suspension was placed on a carbon disk attached to an aluminum sample platform (12.7 mm in diameter) and placed in the oven at 70°C for 15 min until the samples were completely dried. The samples were then analyzed by Scanning Electron Microscopy combined with Energy Dispersive X-ray Spectroscopy (SEM-EDS) using a ProX X-ray spectroscope (Phenom).

### Transmission Electron Microscopy analysis

For Transmission Electron Microscopy (TEM) analysis, cells were prefixed with 1% glutaraldehyde in 10 mM sodium cacodylate buffer (pH 7.2). The fixative was added directly to the growth medium at 1:1 ratio. After 2-h incubation at RT, two washing steps with 10 mM cacodylate buffer were applied. Cells were then immobilized in 2% agarose, post-fixed with 1% osmium tetroxide and 1.5% potassium ferrocyanide, exposed to 1% aqueous thiocarbohydrazide, 2% osmium tetroxide followed by *en bloc* staining with 1% aqueous uranyl acetate and 0.66% lead aspartate. Cells were dehydrated by applying increasing ethanol concentrations (30−100% range), then embedded in epoxy resin, and cut into thin sections (65 nm) using an Ultracut R ultramicrotome (Leica Microsystems, Vienna, Austria). Grids with sections were examined with a JEM 1400 (JEOL Co., Tokyo, Japan, 2008) TEM microscope, equipped with an 11-megapixel TEM camera MORADA G2 (EMSIS GmbH, Münster, Germany).

### Statistical analysis

All measurements were performed in two biological replicas and are reported as mean values ± SD. In the case using technical replicas for each biological replica, this fact is individually specified in each experimental section. ANOVA test (Analysis of Variance) and post-hoc Siegel Tukey test were performed using RStudio Team, 2015. Values with p < 0.05 were considered significantly different. All the graphs were prepared using an OriginPro software (OriginLab Corporation 2018).

## Supplementary data

The following supplemental materials are available:

- Figure S1. Inhibition concentration (IC_50_) of Ni at which *C. merolae* growth is inhibited by 50% after 15 days of exposure.

- Figure S2. Maximal oxygen consumption rates in *C. merolae* cells exposed to 2,4-dinitrophenol (DNP) after 7 days of exposure to 3- and 6-mM Ni.

- Figure S3. Ion maps of Ni^2+^ (red), C_3_H_8_N^+^ (green), Si^+^ (blue) ions overlay over the sample surface in the control (A) and the samples in which *C. merolae* cells were grown in the presence of Ni ions at a concentration of 1 mM (B), 3 mM (C), or 6 mM (D).

- Figure S4. Mass spectral regions for ions selected for the preparation of ion distribution maps.

- Table S1. Quantification of Chl *a* and total carotenoids (pg cell-1) in *C. merolae* cells long-term adapted to various Ni concentrations at different timepoints.

## ACKNOWLEDGEMENTS

This work was supported by the Polish National Science Centre (OPUS 17 grant no. 2019/33/B/NZ3/01870 to J.K. and MINIATURA 5 grant no. 2021/05/X/NZ2/01516 to S.S.). Transmission Electron Microscopy (TEM) experiments were performed at the Nencki Institute, PAS, using infrastructure of the Polish Euro-BioImaging Node. Part of the TEM work was supported by the project financed by the Minister of Education and Science based on contract No. 2022/WK/05 (Polish Euro-BioImaging Node “Advanced Light Microscopy Node Poland”).

